# Chronic activation of a negative engram induces behavioral and cellular abnormalities

**DOI:** 10.1101/2023.11.15.567268

**Authors:** Alexandra L. Jellinger, Rebecca L. Suthard, Bingbing Yuan, Michelle Surets, Evan A. Ruesch, Albit J. Caban, Shawn Liu, Monika Shpokayte, Steve Ramirez

## Abstract

Negative memories engage a brain and body-wide stress response in humans that can alter cognition and behavior. Prolonged stress responses induce maladaptive cellular, circuit, and systems-level changes that can lead to pathological brain states and corresponding disorders in which mood and memory are affected. However, it’s unclear if repeated activation of cells processing negative memories induces similar phenotypes in mice. In this study, we used an activity-dependent tagging method to access neuronal ensembles and assess their molecular characteristics. Sequencing memory engrams in mice revealed that positive (male-to-female exposure) and negative (foot shock) cells upregulated genes linked to anti- and pro-inflammatory responses, respectively. To investigate the impact of persistent activation of negative engrams, we chemogenetically activated them in the ventral hippocampus over three months and conducted anxiety and memory-related tests. Negative engram activation increased anxiety behaviors in both 6- and 14-month-old mice, reduced spatial working memory in older mice, impaired fear extinction in younger mice, and heightened fear generalization in both age groups. Immunohistochemistry revealed changes in microglial and astrocytic structure and number in the hippocampus. In summary, repeated activation of negative memories induces lasting cellular and behavioral abnormalities in mice, offering insights into the negative effects of chronic negative thinking-like behaviors on human health.

## Introduction

Stress has profound impacts on both physical and mental health. In particular, chronic stress is recognized as a potent risk factor for numerous neuropsychiatric disorders, including major depressive disorder (MDD), generalized anxiety disorder (GAD), and post-traumatic stress disorder (PTSD). In humans, repeated negative thinking can parallel the effects of chronic stress. These effects include hippocampal hyperactivity and accelerating the onset of mild cognitive impairment and dementias (Marchant et al., 2020). Additionally, the ability of memories to influence cognition and behavior has been well established. For example, our lab previously demonstrated that reactivating ensembles of hippocampal neurons encoding a positive memory is sufficient to induce anti-depressant effects in mice (Ramirez et al., 2015), and studies in rodents have shown that repeated reactivation of negative memories enhances fear responses (Chen et al., 2019). However, it remains unknown whether modulating a chronic negative memory is sufficient to induce aberrant cellular and behavioral states that recapitulate chronic stress.

The ventral hippocampus (vHPC), specifically vCA1, has been extensively studied for its role in emotional processing. As a modulator of stress responses, vCA1 dysfunction is intimately associated with pathologic affective states (Fanselow and Dong, 2010), and its structure and function is heavily impacted by stressors. Additionally, vCA1 neurons project to various downstream targets and route target-specific streams of information to the basolateral amygdala, nucleus accumbens, and prefrontal cortex (Ciocchi et al., 2015). These diverse connections are integral to the formation of emotional memory, as well as regulating anxiety and the stress response. Within vCA1, stimulation of negative engram cells has been shown to modulate social avoidance behavior and enhance fear responses, but whether chronic stimulation of fearful memories produces enduring effects on brain activity and behavior remains unknown (Chen et al., 2019; Zhang et al., 2019).

Young and old individuals are differentially susceptible to the hippocampal effects of stress and chronic negative thinking behaviors. Epidemiologic population-based studies in humans have established adverse childhood experiences as a risk factor for the development of psychopathologies later in life. Notably, the development of depression and PTSD may increase with the number of these adverse events, suggesting an increased susceptibility of the developing, young brain to the perturbations induced by such events (McLaughlin et al., 2010). However, work in older humans and rodents has shown that there are changes in their stress response and cortisol levels following adverse experiences (Seeman and Robbins, 1994; Sapolsky, Krey and McEwen, 2002). Aged rodents have increased stress vulnerability, as repeated stress increased anxiety-like behaviors, induced spatial memory impairments and raised corticosterone levels more compared to younger animals (Shoji and Mizoguchi, 2010; Wang et al., 2020).

Here, we chronically reactivated negative memory engrams to mimic the effects of stress or chronic negative thinking-related behaviors in mice. We first characterized the genetic landscape of negative and positive memory engrams to search for putative genetic markers involved in healthy and dysfunctional brain states. We then tested whether stimulating negative memory-bearing neurons would be sufficient to induce cellular abnormalities as well as behavioral impairments in young (6-months) and aged (14-months) mice. Our manipulation induced behavioral abnormalities that manifested as increased anxiety, decreased spatial working memory, and increased fear response during extinction and generalization. We also observed cellular alterations, including changes in the number and morphology of microglia and astrocytes. Together, our data suggest that negative memory engrams are a potent means by which to induce cellular and behavioral changes that provide insight into pathological states.

## Methods

### Generation of Single Cell Suspension from Mouse Hippocampal Tissue

As shown in Figure 1 (RNA-sequencing), C57BL/6J mice were bilaterally injected with a 1:1 viral cocktail of AAV9-c-Fos-tTA and AAV9-TRE-eYFP into the ventral hippocampus (vHPC). This activity-dependent labeling strategy captures sufficiently active neurons expressing the immediate-early gene, *cfos*. This system couples the *cfos* promoter to the expression of the tetracycline transactivator (tTA), which binds to the tetracycline response element (TRE) in its protein form. When doxycycline (DOX) is present, such as in the animal’s diet, this inhibits the binding of tTA to TRE, preventing eYFP labeling of active cells (Ramirez et al., 2012; 2014).

**Figure 1:**
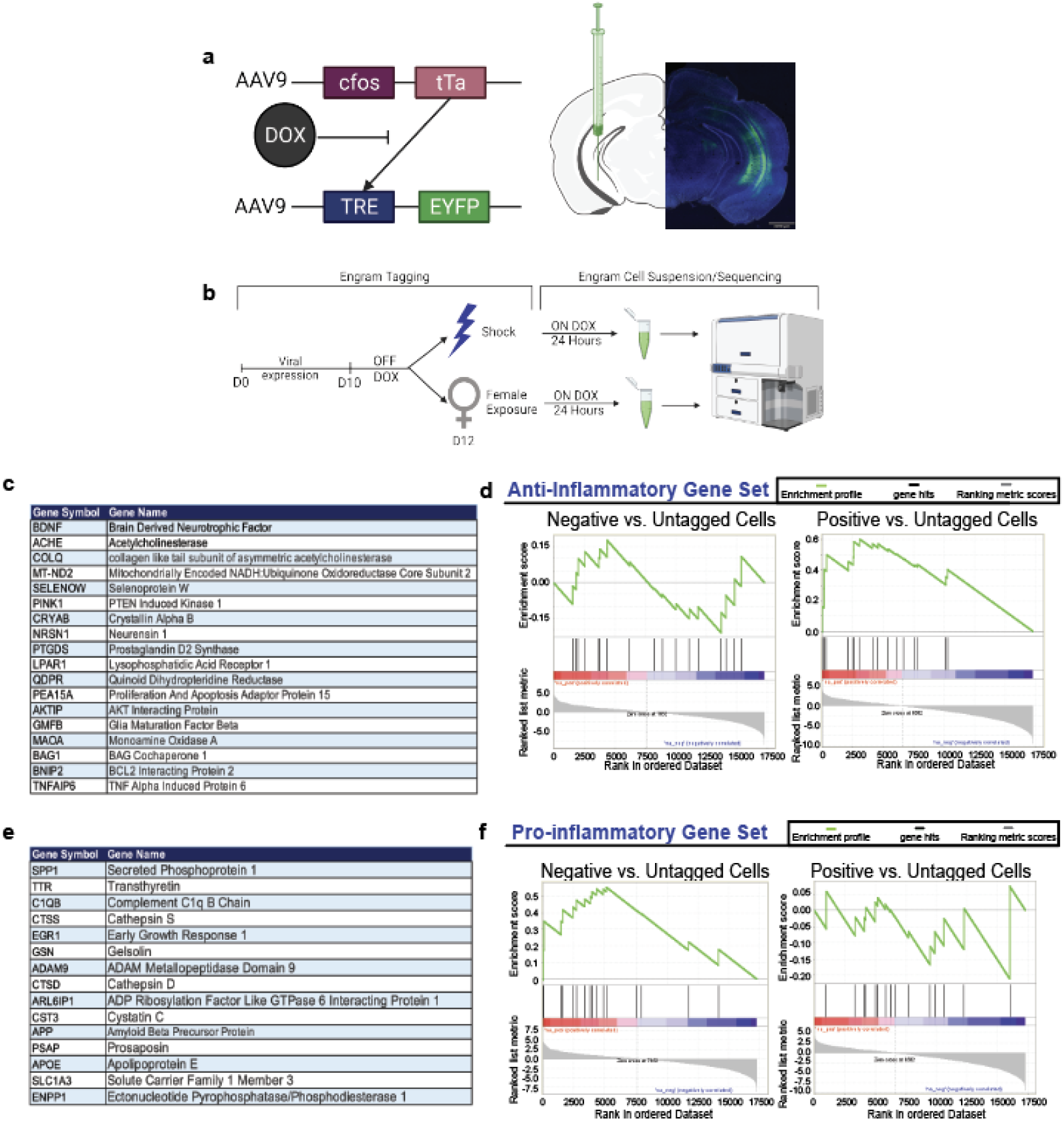
GSEA analysis of RNA-Sequencing data of positive and negative vHPC engram upregulate genes associated with neuroprotective and neurodegeneration respectively. (a) C57BL/6J mice were bilaterally injected with a 1:1 viral cocktail of AAV9-c-Fos-tTA and AAV9-TRE-eYFP into the ventral hippocampus (vHPC). This activity-dependent labeling strategy captures sufficiently active neurons expressing the immediate-early gene, *cfos*. This system couples the *cfos* promoter to the expression of the tetracycline transactivator (tTA), which binds to the tetracycline response element (TRE) in its protein form. When doxycycline (DOX) is present, such as in the animal’s diet, this inhibits the binding of tTA to TRE, preventing eYFP labeling of active cells. (b) Experimental schematic used to label, isolate, and analyze the two groups of vHPC cells labeled by eYFP upon shock (negative) or male-to-female interaction (positive). Prior to surgery, mice were placed on a Dox diet to inhibit the labeling of active neurons. On Day 0, mice were injected with the activity-dependent viral strategy to enable labeling of active neurons during a salient experience. On Day 10 after viral expression and recovery, the mice were taken off of Dox prior to engram tagging to open the labeling window. On Day 11, mice were subjected to contextual fear conditioning (CFC) or male-to-female interaction to label negative or positive neurons, respectively. Mice were placed immediately back on Dox to close this tagging window. 24 hours later, mice were sacrificed and brains were obtained for sequencing experiments. (c) List of gene abbreviations and full names used for the gene set enrichment analysis (GSEA) of anti-inflammatory genes. (d) GSEA analysis with the enrichment score for negative vs. untagged cells and positive vs. untagged cells using the anti-inflammatory gene set. (e) List of gene abbreviations and names used for the GSEA analysis of pro-inflammatory genes. (f) GSEA analysis with the enrichment score of negative vs. untagged cells and positive vs. untagged cells using the pro-inflammatory gene set.

Prior to surgery, mice were placed on a Dox diet to inhibit the labeling of active neurons. After surgery and viral expression, on Day 10, mice were taken off of Dox prior to engram tagging to open the labeling window. On Day 11, mice were subjected to contextual fear conditioning (negative) or male-to-female interaction (positive) to label active neurons with eYFP. Mice were placed immediately back on Dox to close this tagging window. 24 hours later, mice were sacrificed and brains were rapidly extracted, and the hippocampal regions were isolated by microdissection. Eight mice were pooled by each experimental condition. Single cell suspension was prepared according to the guideline of the Adult Brain Dissociation Kit (Miltenyi Botec, Cat No: 13-107-677). Briefly, the hippocampal samples were incubated with digestion enzymes in the C Tube placed on the gentleMACS Octo Dissociator with Heaters with gentleMACS Program: 37C_ABDK_01. After termination of the program, the samples were applied through a MACS SmartStrainer (70 μm). Then a debris removal step and a red blood cell removal step were applied to obtain single cell suspension. Viral injection surgery through RNA-sequencing was performed in Shpokayte et. al., 2022 and this dataset was utilized for further analysis in the current manuscript.

### Isolation of EYFP-positive single cell by FACS

The single cell suspension was subject to a BD FACSAria cell sorter according to the manufacturer’s protocol to isolate eYFP-single cell population. This was performed in Shpokayte et. al., 2022.

### Preparation of RNA-seq library

The RNA of FACS isolated eYFP-positive cells was extracted by using TRIzol (Invitrogen Life Technologies, MA, USA) followed by Direct-zol kit (Zymo Research, CA, USA) according to manufacturer’s instructions. Then the RNA-seq library was prepared using SMART-Seq® v4 Ultra® Low Input RNA Kit (TaKaRa Bio, USA). This was performed in Shpokayte et. al., 2022.

### Analysis of RNA-seq data

The resulting reads from Illumina had good quality by checking with FastQC The first 40 bp of the reads were mapped to mouse genome (mm10) using STAR (Dobin et al., 2013), which was indexed with Ensembl GRCm38.91 gene annotation. The read counts were obtained using featureCounts (Liao, Smyth and Shi, 2014) function from the Subread package with an unstranded option. Reads were normalized by library size and differential expression analysis based on negative binomial distribution was done with DESeq2 (Love, Huber and Anders, 2014). Genes with FDR-adjusted p-value less than 0.00001 plus more than 4-fold difference were considered to be differentially expressed. Raw data along with gene expression levels are being deposited to NCBI Gene Expression Omnibus. For Gene Ontology analysis, we ranked protein-coding genes by their fold changes and used them as the input for the GseaPreranked tool. Based on the GSEA outputs, the dot plots were created with a custom R script. The pathways with FDR-adjusted p-value less than 0.25 were considered to be enriched. GeneMANIA networks were analyzed by GeneMANIA (Warde-Fraley et al., 2010). The max resultant genes were 20 and the max resultant attributes were 10. All networks were assigned an equal weight. The network plots were generated by Cytoscape (Shannon et al., 2003). This was performed in Shpokayte et. al., 2022.

### GSEA Analysis

To further analyze the dataset generated in Shpokayte et al., 2022, we performed further analyses using Gene Set Enrichment Analysis (https://www.gsea-msigdb.org/gsea/index.jsp; Subramanian et al., 2005; Mootha et al., 2003). This is a computational method that determines whether a priori defined set of genes shows statistically significant, concordant differences between two biological states. GSEA is a joint project of UC San Diego and Broad Institute. Gene sets were selected based on existing literature around genes important in neurodegeneration, inflammation and apoptosis.

### Subjects (Figures 2-5)

Fos-based transgenic mouse, Fos-tm2.1(iCre-ERT2) raised on a C57BL/6 background (TRAP2), (Jackson Laboratories; strain #030323) mice were bred in-house. The mice were housed in groups of 2-5 mice per cage. The animal facilities (vivarium and behavioral testing rooms) were maintained on a 12:12-hour light cycle (0700-1900). Mice received food and water *ad libitum* before surgery. Following surgery, mice were group-housed with littermates and allowed to recover for a minimum of 4 weeks before experimentation for viral expression. All subjects were treated in accord with protocol 201800579 approved by the Institutional Animal Care and Use Committee (IACUC) at Boston University.

**Figure 2:**
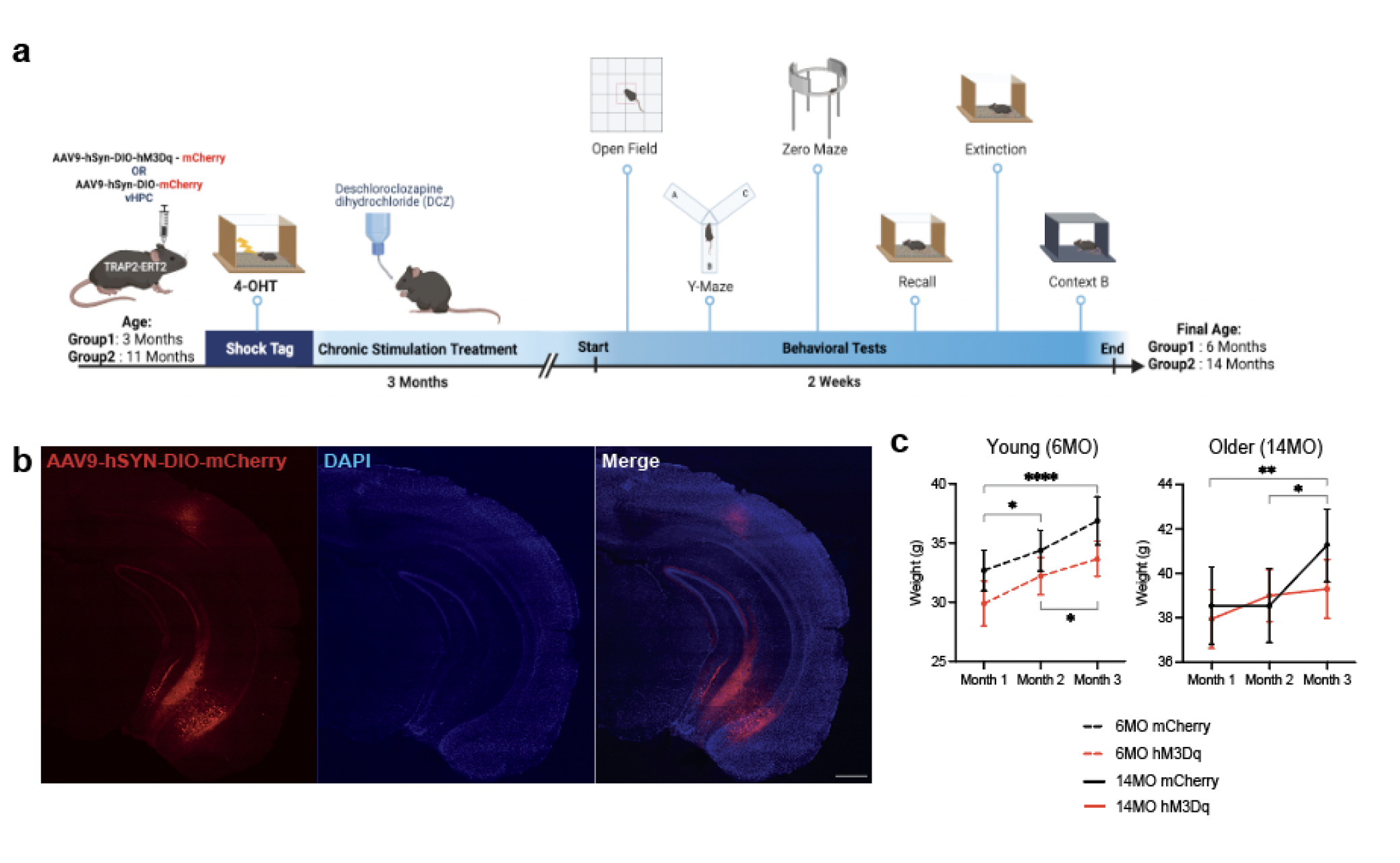
Experimental timeline for chronic modulation of a negative engram in TRAP2 mice. (a) Schematic representation of tagging a negative engram in young (3 month) and old (11 month) TRAP2 mice followed by three months of chronic engram stimulation and a battery of behavioral tests. AAV9-hSyn-DIO-hM3Dq-mCherry or AVV9-hSyn-DIO-mCherry control vector was bilaterally injected into the vHPC of young and old mice. After viral expression and surgical recovery, all mice were subjected to contextual fear conditioning (CFC) in Context A and subsequent 4-hydroxytamoxifen (4-OHT) intraperitoneal (IP) injection to induce negative engram tagging. For the next three months, mice received the water-soluble Designer Receptors Exclusively Activated by Designer Drugs (DREADD) agonist deschloroclozapine dihydrochloride (DCZ) in their home cage water. After three months of stimulation wherein the young and old groups reached 6 months and 14 months of age, the mice underwent open field, y-maze, zero maze, remote recall, extinction, and generalization in a novel Context B. (b) Representative image of hSyn-DIO-hM3Dq-mCherry expression in the vHPC (red) and DAPI+ cells (blue) after 3 months of hM3Dq activation. (c) Weights of all groups were recorded over the three month stimulation protocol, once per month. (left) young 6-month old and (right) older 14-month old mice. Values are given as a mean + (SEM). Statistical analysis was performed with two-way (RM) ANOVA followed by Sidak’s post-hoc test. n=9-17 per group after outlier removal. p ≤ 0.05, **p ≤ 0.01, ***p ≤ 0.001, ****p ≤ 0.0001, no label = not significant

### DCZ administration

All animals had their home cage water supply replaced with water-soluble deschloroclozapine dihydrochloride (DCZ) in distilled water (diH2O) every 5 days (HelloBio, #HB9126). Administration was calculated based on each mouse drinking approximately 6mL of water per day. A stock solution was created of 1-2mg of water soluble DCZ into 1 mL of DMSO and vortexed until in solution. 17ul of stock solution was combined with 1L of diH20 for a final concentration of 3ug/kg/day per mouse, as previously described in Suthard et. al., 2023, to avoid any potential excitotoxicity resulting through stimulating cells for up to 3 months, and to mimic a more sustained dysregulation that may accumulate in severity over time. Importantly, no obvious signs of distress or seizures were observed with chronic administration of DCZ by experimenters or vivarium staff for the entire duration of the project.

### Stereotaxic surgeries

For all surgeries, mice were initially anesthetized with 3.0% isoflurane inhalation during induction and maintained at 1-2% isoflurane inhalation through stereotaxic nose-cone delivery (oxygen 1 L/min). Ophthalmic ointment was applied to the eyes to provide adequate lubrication and prevent corneal desiccation. The hair on the scalp above the surgical site was removed using Veet hair removal cream and subsequently cleaned with alternating applications of betadine solution and 70% ethanol. 2.0% lidocaine hydrochloride (HCl) was injected subcutaneously as local analgesia prior to midsagittal incision of the scalp skin to expose the skull. 0.1mg/kg (5mg/kg) subcutaneous (SQ) dose of meloxicam was administered at the beginning of surgery. All animals received bilateral craniotomies with a 0.5-0.6 mm drill-bit for vCA1 injections. A 10uL airtight Hamilton syringe with an attached 33-gauge beveled needle was slowly lowered to the coordinates of vCA1: -3.16 anteroposterior (AP), ± 3.10 mediolateral (ML) and both -4.25/-4.50 dorsoventral (DV) to cover vertical the axis of ventral hippocampus. All coordinates are given relative to bregma (mm). A total volume of 300nL of AAV9-DIO-Flex-hM3Dq-mCherry or AAV9-DIO-Flex-mCherry was bilaterally injected into the vCA1 at both DV coordinates (150nL at each site) listed above. The needle remained at the target site for 7-10 minutes post-injection before removal. Incisions were sutured closed using 4/0 Non-absorbable Nylon Monofilament Suture [Brosan]. Following surgery, mice were injected with 0.1mg/kg intraperitoneal (IP) dose of buprenorphine (volume administered was dependent on the weight of the animal at the time of surgery). They were placed in a recovery cage with a heating pad until fully recovered from anesthesia. Histological assessment verified bilateral viral targeting and data from off-target injections were not included in analyses.

### Immunohistochemistry

Mice were overdosed with 3% isoflurane and perfused transcardially with cold (4C) phosphate-buffered saline (PBS) followed by 4% paraformaldehyde (PFA) in PBS. Brains were extracted and kept in PFA at 4C for 24-48 hours and transferred to a 30% sucrose in PBS solution. Long-term storage of the brains consisted of transferring to a 0.01% sodium azide in PBS solution until slicing. Brains were sectioned into 50 um thick coronal and sagittal sections with a vibratome and collected in cold PBS or 0.01% sodium azide in PBS for long-term storage. Sections underwent three washes for 10 minutes each in PBS to remove the 0.01% sodium azide that they were stored in. Sections were blocked for 2 hours at room temperature (RT) in PBS combined with 0.2% Triton X-100 (PBST) and 5% normal goat serum (NGS) on a shaker. Sections were incubated in the primary antibody (1:500 polyclonal guinea pig anti-NeuN/Fox3 [SySy #266-004]; 1:1000 mouse anti-GFAP [NeuroMab #N206A/8]; 1:1000 rabbit polyclonal anti-Iba1 [Wako #019-19741]) diluted in the same PBST-NGS solution at 4°C for 24-48 hours. Sections then underwent three washes for 10 minutes each in PBST. Sections were incubated in a secondary antibody (1:200 Alexa Fluor 488 anti-guinea [Invitrogen #A-11073], 1:1000 Alexa Fluor 488 anti-rabbit [Invitrogen #A-11008], 1:1000 Alexa Fluor 555 anti-mouse [Invitrogen #A-11001]) for 2 hours at RT. Sections then underwent three more washes in PBST for 10 minutes each. Sections were then mounted onto microscope slides (VMR International LLC, PA, USA). Vectashield HardSet Mounting Medium with DAPI (Vector Laboratories Inc, CA, USA) was applied and slides were coverslipped and allowed to dry for 24 hours at RT. Once dry, slides were sealed with clear nail polish around the edges and stored in a slide box in 4°C. If not mounted immediately, sections were stored in PBS at 4°C.

### Cell Quantification and Morphological Analysis (NeuN, Iba1, GFAP)

Only animals that had accurate bilateral injections were selected for quantification. Images were acquired using a confocal microscope (Zeiss LSM800, Germany) with a 20X objective. 6-8 single tile images were taken in the brain regions of interest across 3-5 coronal slices giving us a total of 18 single tile images per mouse x 3-4 mice in each group. When imaging NeuN z-stacks, intervals were taken at 4um, whereas Iba1 and GFAP z-stacks were taken at 1um intervals for detailed morphological analysis. Ilastik (Berg et. al., 2019) was used to quantify the total number of cells and 3DMorph (York et. al., 2018) was used to assess the morphology of GFAP and Iba1 cells. Images of NeuN, GFAP, and Iba-1 were captured in a 300 x 300 single-tile (micrometers) that contained matched ROIs across groups for hippocampal subregions.

For 3DMorph analysis, we selected a total of 3 single tile images from the same ROI per mouse per group. This gave a total of ∼20-30 cells per image. Therefore, we conducted morphological analysis on a total of 755 Iba1+ cells and 842 GFAP+ cells across all groups. Acquired image files (.czi) were opened in ImageJ. Processing of images in Figures 1, 2, 5 and Supplemental 1 involved maximizing intensity, removing outlier noise (despeckling), and adjusting contrast of images for improved detection of glial cells in 3DMorph.

Cell volume (um^3^) is calculated by converting the number of voxels occupied by the cell object to a real-world unit based on defined scaling factors. Territorial volume (um^3^) is estimated by creating a polygon around the outside points of each cell and measuring the volume inside. Ramification index is a measure of the complexity of astrocyte and microglial morphology, with a higher ramification index indicating a more complex and extensive branching structure. This is computed by dividing the cell territorial volume by the cell volume. Average centroid distance (um) helps to understand the distribution of cells in the brain, with a larger centroid distance indicating the glial cells are further apart. Using a 3D reconstruction of each cell skeleton, branches are labeled as primary, secondary, tertiary and quaternary. The number of branch points (#) are calculated by the intersections of each branch type and endpoints (#) are counted at the termination of these branches. Finally, each of these branches are measured in length and utilized for the minimum, maximum and average branch length (um).

### Behavioral assays

All behavior assays were conducted during the light cycle of the day (0700-1900) on animals. Mice were handled for 3–5 days, 2 min per day, before all behavioral experiments began. Behavioral assays include open field, zero maze, y-maze, fear conditioning, recall, extinction, and generalization. As described in our previous work in greater detail (Suthard et. al., 2023), DCZ water was removed 1-2 hours from the home cage to allow for a decline in ligand concentration in the brain and to ensure there were no acute effects of the drug on behavior. When administered intraperitoneally (I.P.) in mice, DCZ peaks 5-10 minutes after injection, declining to baseline levels around 2 hours after (Nagai et. al., 2020).

### Open field test

An open (61cm x 61cm) arena with black plastic walls was used for the open field test, with a red-taped area in the middle delineating “center” from “edges”. A camera was placed above the open field in order to record video of the session. Mice were individually placed into the center of the chamber, and allowed to explore freely for 10 min. At the end of the session, each mouse was placed into a separate cage until all of its cage mates had also gone through the behavioral test. They were all placed back into the home cage once this occurred. An automated video-tracking system, AnyMaze v4.3 (Stoelting Co., Wood Dale, Il) was used to measure total distance traveled, number of entries into the center, and total time spent in the center.

### Zero maze test

The zero maze test is a pharmacologically-validated assay of anxiety in animal models that is based on the natural aversion of mice to elevated, open areas. It is composed of a 44cm wide ring with an outer diameter of 62cm. It contained four equal zones of walled (closed) and unwalled (open) areas. The entire ring is 8 cm in height and sits at an elevation of 66cm off the ground. All animals at each timepoint were tested on the same day. Mice were placed in the closed area at the start of a 10-minute session. The following parameters were analyzed: time spent in the open area, number of entries into the open area, etc. The maze was cleaned with 70% ethanol between mice. During the behavioral testing, the lighting levels remained constant and there were no shadows over the surface of the maze.

### Y-maze test

The Y-maze is a hippocampal-dependent spatial working memory task that requires mice to use external cues to navigate the identical internal arms. The apparatus consisted of a clear plexiglass maze with three arms (36.5cm length, 7.5cm width, 12.5cm height) that were intersected at 120 degrees. A mouse was placed at the end of one arm and allowed to move freely through the maze for 10 minutes without any reinforcements, such as food or water. Entries into all arms were noted (center of the body must cross into the arm for a valid entry) and a spontaneous alternation was counted if an animal entered three different arms consecutively. Percentage of spontaneous alternation was calculated according to the following formula: [(number of alterations)/(total number of arm entries - 1) x 100]. To prevent bias in data analysis, the test was carried out in a blind manner by the experimenter and behavior was analyzed blindly in AnyMaze. Additionally, total distance traveled was calculated for the test to rule out changes in locomotion.

### Fear conditioning, recall and extinction

For contextual fear conditioning (CFC) for all timepoints took place in mouse conditioning chambers (Coulbourn Instruments, Holliston, MA). The grid floor was connected to a precision animal shocker (Coulbourn Instruments, Holliston, MA) which delivered a total of four foot shocks (2s duration, 1.5mA) at the 120, 180, 240 and 300 second time points during the 360 second session. This mouse conditioning chamber (18.5 x 18.5 x 21.5cm) was composed of metal walls, plexiglass front and back walls and a stainless-steel grid floor (16 grid bars). For recall, all animals were placed back into the same chambers they were originally fear conditioned in for a total of 300 seconds in the absence of foot shocks. For extinction, all animals were placed back into the same chambers they were originally conditioned in for a total of 1800 seconds in the absence of foot shocks. To record the animal’s freezing activity, a video camera was mounted to the ceiling of the chamber and fed into a computer running FreezeFrame software (Actimetrics, Wilmette, IL). The program determined movement as changes in pixel luminance over a set period of time. Freezing was defined as a bout of 1.25 s or longer without changes in pixel luminance and verified by an experimenter blind to treatment groups. The chambers were cleaned with 70% ethanol solution prior to each animal placement.

### Context B Exposure

For generalization (Context B), after 5 days of extinction, the animals were taken into a different room from the original conditioning and placed in a new larger chamber. During the experiment the floor was changed to a smooth bottom, the walls were changed to have striped backgrounds, the odor was changed to a neutral almond scent, and the lights were turned off during the procedure. Red lights were turned on in the room for the experimenter. The entire exposure to context B was a total of 10 minutes. Activity recording and freezing level analysis was assessed as described above using FreezeFrame Software. The chambers were cleaned with 70% ethanol solution prior to each animal placement.

### Statistics & Data Analysis

Data were analyzed using GraphPad Prism (GraphPad Software v9.4.1, La Jolla, CA). Data were tested for normality using the Shapiro-Wilk and Kolmogorov-Smirnov tests. Outliers were removed using the ROUT method, which uses identification from nonlinear regression. We chose a ROUT coefficient Q value of 10% (False Discovery Rate), making the threshold for outliers less-strict and allowing for an increase in power for outlier detection. No more than 1-2 mice were removed in any group as outliers for any behavioral measure. To analyze differences between groups (mCherry and hM3Dq) and across age groups (3- and 11-months old [Fear Conditioning]; 6- and 14-months old [All other behavioral tasks]) we used a two-way ANOVA (between-subject factor: Group; within-subject factor: Age). To analyze differences between groups and across time within a single session (i.e. fear conditioning) or across months (i.e. body weight), we used two-way repeated measures (RM) ANOVAs (between-subjects factor: Group; within-subjects factor: Time/Month). Post-hoc analyses were performed using Tukey’s and Sidak’s multiple comparisons test, where applicable. Alpha was set to 0.05 for all tests.

## Results

### Ventral hippocampal negative and positive engram cells differentially express genes associated with inflammation

The ventral hippocampus (vHPC) parses out negative and positive memories into anatomically segregated, functionally distinct, and molecularly diverse neuronal ensembles or ‘engrams’ that have unique transcriptional profiles and DNA methylation landscapes (Shpokayte et. al., 2022). As positive and negative memories have been respectively implicated in driving approach and avoidance-related behaviors in mice, we first sought to test if these two vHPC engrams differentially upregulated genes in response to each experience. We previously performed a RNA-seq experiment described in Shpokayte et. al., 2022. Briefly, we utilized an activity-dependent labeling technique to visualize and manipulate the cells processing a specific memory in the hippocampus by leveraging the cFos protein as an indicator of neuronal activity (Liu & Ramirez et. al., 2012). Our approach combines a doxycycline (DOX)-dependent strategy with a dual-viruses approach consisting of AAV9-c-fos-tTA + AAV9-TRE-ChR2-eYFP or control AAV9-TRE-eYFP, 24 hours prior to a behavioral experience, DOX is removed from the animal’s diet which will allow tTa to bind to TRE and drive the expression of any downstream effector, such as the fluorescent protein eYFP (**Figure 1A**). After the behavioral experience, i.e. contextual fear conditioning or male-to-female exposure, the animal is placed back on DOX to close the tagging window for the negative or positive experience. Thus, the cells involved in processing a discrete experience expresses the eYFP protein which can be used in fluorescence-activated cell sorting (FACS) to isolate only the eYFP+ as ‘tagged’ engram cells for genetic sequencing. To analyze this RNA-seq dataset, we selected specific gene sets and conducted Gene Set Enrichment Analysis (GSEA) to compare the up- and down-regulation of these genes in positive or negative engram cells. The GSEA analysis showed that the anti-inflammatory gene set was enriched in positive engram cells, with higher expression compared to negative engram cells (**Figure 1C-D**). Further, the pro-inflammatory gene set was enriched in negative engram cells, with higher expression compared to positive engram cells (**Figure 1E-F**). Interestingly, this gene set included *Spp1*, *Ttr*, and *C1qb1*, which are involved in inflammatory processes and oxidative stress. Together, these data provide the start to uncovering the molecular landscape that positive and negative memory-bearing cells may contain that links them to protective or pathological functioning, respectively.

### Chronic engram stimulation increases anxiety-related behaviors, decreases spatial memory and impairs fear extinction and generalization

Mouse models of chronic stress display increased anxiety-related behaviors and impairments in cognition and memory-related tasks that are similar to those seen in human patients with MDD, PTSD and comorbid GAD (Tran and Gellner, 2023). To further understand the impact of negative engram cells, we next asked if chronic stimulation of a negative engram could induce behavioral changes in anxiety and memory-related tasks. To that end, we combined excitatory Designer Receptors Exclusively Activated by Designer Drugs (DREADDs) with a Fos-based transgenic mouse, Fos-tm2.1(iCre-ERT2) raised on a C57BL/6 background (TRAP2), to label active cells under the control of 4-Hydroxytamoxifen (4-OHT) (DeNardo et. al., 2019). 3- and 11-month-old TRAP2 mice were bilaterally injected with a Cre-dependent AAV-DIO-hM3Dq-mCherry or control AAV-DIO-mCherry into the vHPC. After two weeks for viral expression and surgical recovery, all mice were subjected to CFC, consisting of four, 1.5mA foot shocks in Context A (CxtA). 30 minutes after CFC, mice received a 40 mg/kg injection of 4-OHT to label cells sufficiently active during this fearful experience (**Figure 2A-B**). The animals were returned to their home cages, where they received water-soluble deschloroclozapine dihydrochloride (DCZ), a highly potent, selective, and brain-penetrable muscarinic hM3D(Gq) actuator, for three months in their daily water (Nagai et al., 2020). DCZ was selected because even at very low concentrations, this ligand is able to increase neuronal activity with binding to hM3D(Gq) receptors (Nagai et. al., 2020). To avoid unnecessary pain or stress to the animal with chronic administration of DCZ intraperitoneally (I.P.), we selected the voluntary oral route (P.O.), which has been utilized in the past for chemogenetic experiments (Zhan et. al., 2019 and shows no significant differences in behavior at the same dose (Ferrari et. al., 2022).

After three months of negative engram stimulation, mCherry and hM3Dq mice were aged to 6- and 14-months old at the time of behavioral testing (**Figure 2A**). The weights of both the young and old mice increased independently of hM3Dq activation across the three months of stimulation, suggesting normal weight gain with aging (Two-way RM ANOVA; [6 Months] Interaction: F(2,34) = 0.2633, p=0.7701; Month: F(2,34) = 14.84, p<0.0001; Group: F(1,17) = 1.361, p=0.2594; Subject: F(17,34) = 15.44, p<0.0001. Sidak’s [Month]: 1 vs. 2, p=0.0279; 1 vs. 3, p<0.0001; 2 vs. 3, p=0.0323). [14 Months] Interaction: F(2,52) = 2.607, p=0.0833; Month: F(2,52) = 7.854, p=0.0010; Group: F(1,26) = 0.1275, p=0.7239; Subject: F(26,52) = 20.74, p<0.0001. Sidak’s [Month]: 1 vs. 2, p=0.6942; 1 vs. 3, p=0.0011; 2 vs. 3, p=0.0198)(**Figure 2C**). However, we note that we did not include a Month 0 body weight measurement, thus, it is possible that during the first month of stimulation (Month 0-1), there may have been a drop in body weight that rebounded by the first measurement at Month 1 that continued to increase normally through Months 2-3 as shown in Figure 1.

Importantly, there were no decreases in the number of NeuN+ cells in the subiculum, dentate gyrus and CA1 of the ventral hippocampus due to chronic activation, suggesting that we are not inducing a reduction in neuronal cell health with this constant stimulation, as measured by this method (Two-way ANOVA; [Subiculum] Interaction: F(1,8) = 0.2516, p=0.6295; Age: F(1,8) = 7.341, p=0.0267; Group: F(1,8) = 0.00721, p=0.9344. Tukey’s [Group]: 6MO mCherry vs. hM3Dq, p=0.9904; 14MO mCherry vs. hM3Dq, p=0.9744. (**Supplemental Figure 1A-B**). [vDG] Interaction: F(1,8) = 0.6806, p=0.4333; Age: F(1,8) = 4.423, p=0.0686; Group: F(1,8) = 0.0524, p=0.8246. (**Supplemental Figure 1C-D**). [vCA1] Interaction: F(1,8) = 2.824, p=0.1314; Age: F(1,8) = 0.5443, p=0.4817; Group: F(1,8) = 2.947, p=0.1244)(**Supplemental Figure 1E-F**).

To test the hypothesis that chronic activation of a negative memory is sufficient to increase anxiety-like behaviors, we subjected our mice to the open field and zero maze tests (**Figure 3**). In the open field (**Figure 3A**), 6- and 14-month-old hM3Dq mice spent less time in the center compared to age-matched mCherry mice (Two-way ANOVA; Interaction: F(1,33) = 0.1438, p=0.7070; Age: F(1,33) = 0.5858, p=0.4495; Group: F(1,33) = 29.56, p<0.0001)(Tukey’s multiple comparisons [Group]: 6MO mCherry vs. hM3Dq, p=0.0049; 14MO mCherry vs. hM3Dq, p=0.0016)(**Figure 3A-ii**). We observed no significant differences in total distance traveled across all groups (Two-way ANOVA; Interaction: F(1,37) = 0.0016, p=0.9675; Age: F(1,37) = 0.6862, p=0.4128; Group: F(1,37) = 0.3522, p=0.5565)(**Figure 3A-i**).

**Figure 3:**
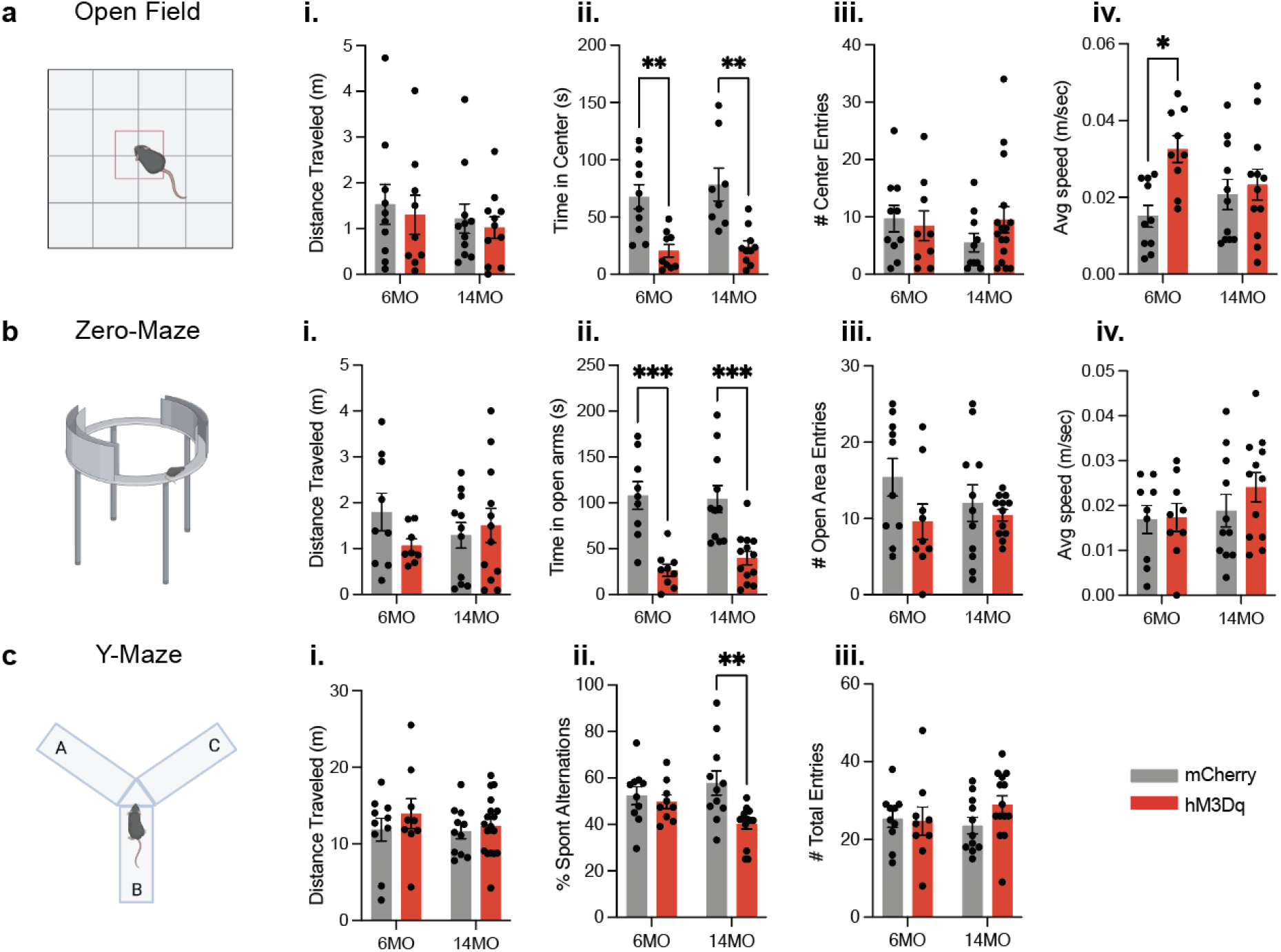
Chronic activation of an aversive engram induces behavioral changes in anxiety and working memory. (a) Schematic of open field test; (i.) total distance traveled, (ii.) total time spent in center, (iii.) number of entries into the center, and (iv.) average speed in 6 month mCherry and hM3Dq and 14 month mCherry and hM3Dq after 3 months of stimulation. (b) Schematic of zero-maze test; (i.) total distance traveled, (ii.) total time spent in open arms, (iii.) number of entries into the open area, and (iv.) average speed. (c) Schematic of Y-maze test; (i.) total distance traveled, (ii.) percentage of spontaneous alterations, and (iii.) total number of entries into arms. Statistical measure used a two-way ANOVA with time point and group as factors. Tukey’s post-hoc tests were performed. Error bars indicate SEM. p ≤ 0.05, **p ≤ 0.01, ***p ≤ 0.001, ****p ≤ 0.0001, ns = not significant. n= 8-11 per group after outlier removal.

Further, 6-month-old hM3Dq mice had an increase in average speed in the open field compared to mCherry mice (Two-way ANOVA; Interaction: F(1,38) = 4.003, p=0.526; Age: F(1,38) =0.2346, p=0.6306; Group: F(1,38) = 7.306, p=0.0102)(Tukey’s [Group]: 6MO mCherry vs. hM3Dq, p=0.0150; 14MO mCherry vs. hM3Dq, p=0.9532)(**Figure 3A-iv**). However, there were no differences in the number of center entries across all groups, indicating that mice in mCherry and hM3Dq groups are entering the center the same number of times, but spending less time per visit (Two-way ANOVA; Interaction: F(1,41) = 1.236, p=0.2728; Age: F(1,41) = 0.4424, p=0.5097; Group: F(1,41) = 0.3370, p=0.5648)(**Figure 3A-iii**).

In the zero-maze (**Figure 3B**), the 6- and 14-month-old hM3Dq mice spent less time in the center compared to age-matched mCherry mice (Two-way ANOVA; Interaction: F(1,38) = 0.5633, p=0.4575; Age: F(1,38) = 0.1709, p=0.6816; Group: F(1,38) = 40.22, p<0.0001)(Tukey’s [Group]: 6MO mCherry vs. hM3Dq, p=0.0002; 14MO mCherry vs. hM3Dq, p=0.0007)(**Figure 3B-ii**). No significant differences were observed in the total distance traveled (**Figure 3B-i**) or number of entries into the open area across all groups (Two-way ANOVA; [Total Distance Traveled] Interaction: F(1,36) = 1.915, p=0.1749; Age: F(1,36) = 0.0093, p=0.9237; Group: F(1,36) = 0.6022, p=0.4428. [Open Area Entries] Interaction: F(1,38) = 1.082, p=0.3049; Age: F(1,38) = 0.3839, p=0.5392; Group: F(1,38) = 3.286, p=0.0778)(**Figure 3B-iii**). These findings indicate that mCherry and hM3Dq mice entered the open area the same number of times, but hM3Dq mice had shorter visits. We also observed no significant difference in average speed across all groups (Two-way ANOVA; Interaction: F(1,37) = 0.5118, p=0.4788; Age: F(1,37) = 1.659, p=0.2057; Group: F(1,37) = 0.7180, p=0.4023)(**Figure 3B-iv**). Ultimately, our observations from open field and zero maze support the hypothesis that chronic activation of a negative memory is sufficient to increase anxiety-like behaviors.

The mice also underwent y-maze testing in order to test the hypothesis that chronic activation of a negative memory is sufficient to impair spatial working memory (**Figure 3C**). In this task, mice were required to remember the most recent arm of the maze that was explored and subsequently navigate to a ‘novel’ arm. Mice freely explored the three arms of the maze (A, B, C) and spatial working memory was assessed by measuring the percentage of spontaneous alternations (i.e. ABC, CBA, CAB), or three consecutive entrances to a novel arm without repetition. A lower percentage of spontaneous alternations indicates a deficit in spatial working memory, as they can not recognize what arm they were in just moments prior. We observed a higher percentage of spontaneous alterations in 14-month-old mCherry than 14-month-old hM3Dq mice (Two-way ANOVA; Interaction: F(1,40) = 4.088, p=0.0499; Age: F(1,40) = 0.3080, p=0.5820; Group: F(1,40) = 7.517, p=0.0091)(Tukey’s [Group]: 14MO mCherry vs. hM3Dq, p=0.0045)(**Figure 3C-ii**). However, no significant difference in the percentage of spontaneous alterations was observed between 6-month-old mCherry and hM3Dq mice (Tukey’s [Group]: 6MO mCherry vs. hM3Dq, p=0.9634)(**Figure 3C-ii**). Additionally, we observed no significant differences in the total distance traveled or total number of entries across all groups (Two-way ANOVA; [Total Distance Traveled] Interaction: F(1,44) = 0.2668, p=0.6081; Age: F(1,44) = 0.5106, p=0.4787; Group: F(1,44) = 1.181, p=0.2831 (**Figure 3C-i**). [Total Number of Entries] Interaction: F(1,40) = 1.332, p=0.2553; Age: F(1,40) = 0.2236, p=0.6389; Group: F(1,40) = 0.8247, p=0.3692 (**Figure 3C-iii**). Therefore, our observations suggest an inability of the mice to remember which arms of the Y-maze they previously explored and may represent an impairment in working memory in the 14-month cohort.

Finally, mice had a ‘negative’ engram tagged in a fearful CxtA before the initiation of chronic activation (**Figure 4A**). During this contextual fear conditioning session, mCherry and hM3Dq mice in the 3- and 11-month groups displayed no differences in freezing behavior, as expected (Two-way ANOVA; Interaction: F(1,42) = 0.1342, p=0.7150; Age: F(1,42) = 12.20, p=0.0011; Group: F(1,42) = 2.913, p=0.0953)(**Figure 4Ai-ii**). However, to test the hypothesis that chronic negative engram activation is sufficient to induce impaired fear memory recall and extinction, mice were subjected to remote memory recall after 3 months of treatment. The only significant difference in remote recall behavior was an increase in freezing between 6 and 14 month mCherry mice (Two-way ANOVA; Interaction: F(1,41) = 2.531, p=0.1193; Age: F(1,41) = 10.08, p=0.0028; Group: F(1,41) = 6.406, p=0.0153)(Tukey’s [Age]: 6 vs. 14MO mCherry, p=0.0101; 6 vs. 14MO hM3Dq, p=0.6672. [Group] 6MO mCherry vs. hM3Dq, p=0.0449; 14MO mCherry vs. hM3Dq, p=0.8890)(**Figure 4Bi-ii**). Mice typically display increased freezing behavior as they age, so these effects during remote recall are expected (Shoji & Miyakawa, 2019). For the extinction session in the same context, there were no significant differences across all groups at 6- and 14-months of age (**Figure 4Ci-ii**). However, we observed a strong trend showing an increase in freezing in the 6-month-old hM3Dq mice that was like remote recall (Two-way ANOVA; Interaction: F(1,41) = 2.618, p=0.1133; Age: F(1,41) = 2.335, p=0.1342; Group: F(1,41) = 5.763, p=0.0210)(Tukey’s [Group]: 6MO mCherry vs. hM3Dq, p=0.0590; 14MO mCherry vs. hM3Dq, p=0.9269)(**Figure 4Di-ii**). This may suggest that these mice are displaying an increase in fear during remote recall of a fearful experience from 3 months prior and have difficulty appropriately decreasing levels of fear during extinction. Finally, mice were placed in a neutral, novel context B at this remote time point after chronic activation to test for generalization of fear. In this context B, mice in the hM3Dq groups at both 6- and 14-months of age displayed increased freezing in a neutral context compared to mCherry controls (Two-way ANOVA; Interaction: F(1,41) = 0.09801, p=0.7558; Age: F(1,41) = 2.624, p=0.1129; Group: F(1,41) = 26.53, p<0.0001)(Tukey’s [Group]: 6MO mCherry vs. hM3Dq, p=0.0053; 14MO mCherry vs. hM3Dq, p=0.0026)(**Figure 4G-H**). Chronic negative engram activation induced selective deficits in extinction and modestly in fear generalization, which may point to the negative impact of fearful memories in human patients with disorders like PTSD (Maren and Holmes, 2016).

**Figure 4:**
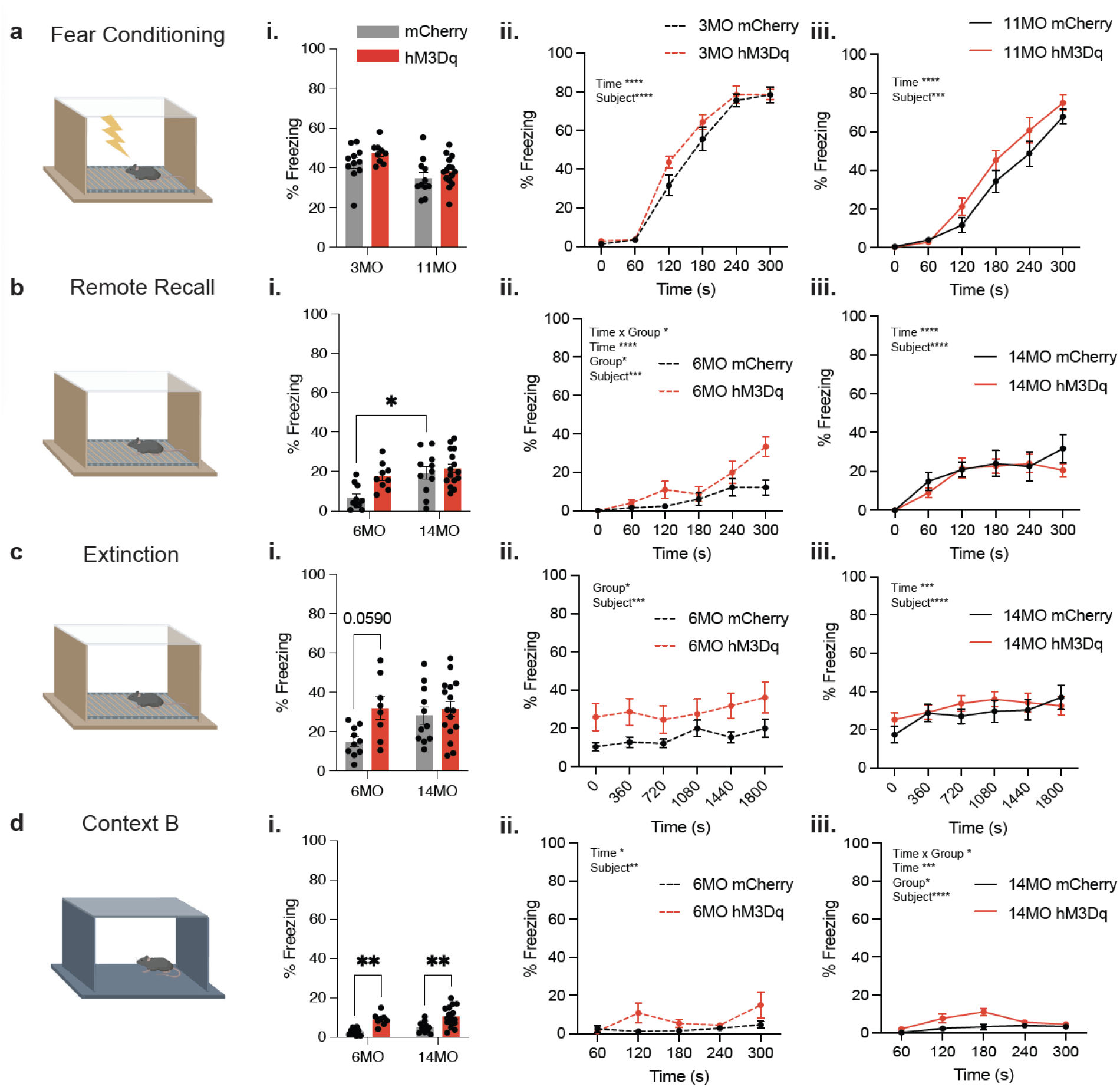
Chronic activation of the negative engram induces behavioral changes in fear memory. (a) Schematic of contextual fear conditioning (CFC) in context A wherein 3- and 11-month-old mice received four, 1.5mA, 2 second foot shocks. Following the CFC session, the negative engram was tagged in all mice by IP injection of 4-OHT. (i.) Average percent freezing levels and total percentage freezing across the 300 second CFC session for (ii.) 6 month groups and (iii.) 14 month groups [Two-way ANOVA RM; [6 Months] Interaction: F(5,130) = 1.131, p=0.3473; Time: F(5,130) = 115.6, p<0.0001; Group: F(1,26) = 2.737, p=0.1101; Subject: F(26,130) =3.056, p<0.0001. [14 Months] Interaction: F(5,90) = 1.218, p=0.3074; Time: F(5,90) = 234.4, p<0.0001; Group: F(1,18) = 1.862, p=0.1892; Subject: F(18,90) = 2.887, p=0.0005]. (b) Schematic of remote recall performed after 3 months of negative engram stimulation. Mice were returned to context A for 5 minutes in the absence of foot shocks. (i.) Average percent freezing levels and total percentage freezing across the 300 second remote recall session for (ii.) 6 month groups and (iii.) 14 month groups [Two-way ANOVA RM; [6 Months] Interaction: F(5,85) = 15.72, p=0.0115; Time: F(5,85) = 15.72, p<0.001; Group: F(1,17) = 6.588, p=0.0200; Subject: F(17,85) = 2.458, p=0.0035. [14 Months] Interaction: F(5,130) = 0.9027, p=0.4815; Time: F(5,130) = 14.18, p<0.0001; Group: F(1,26) = 0.3967, p=0.5343; Subject: F(26,130) = 4.107, p<0.0001]. (c) Schematic of extinction wherein mice were placed in context A for 30 minutes in the absence of foot shocks. (i.) Average percent freezing levels and total percentage freezing across the 1800 second extinction session for (ii.) 6 month groups and (iii.) 14 month groups [Two-way ANOVA RM; [6 Months] Interaction: F(5,85) = 0.4133, p=0.8383; Time: F(5,85) = 1.979, p=0.0899; Group: F(1,17) = 5.494, p=0.0315; Subject: F(17,85) = 7.352, p<0.0001. [14 Months] Interaction: F(5,130) = 1,362, p=0.2570; Time: F(5,130) = 5.387, p=0.0002; Group: F(1,26) = 0.3836, p=0.5411; Subject: F(26, 130) = 11.16, p<0.0001]. (d) Schematic of the novel context B used to assess generalization. (i.) Average percent freezing levels and total percentage freezing across the 300 second generalization session for (ii.) 6 month groups and (iii.) 14 month groups [Two-way ANOVA RM; [6 months] Interaction: F(4,72) = 2.257, p=0.0713; Time: F(4,72) = 3.373, p=0.0138; Group: F(1,18) = 4.151, p=0.0566; Subject: F(18,72) = 2.537, p=0.0028. [14 Months] Interaction: F(4,100) = 2.575, p=0.0421; Time: F(4,100) = 5.717, p=0.0003; Group: F(1,25) = 6.343, p=0.0186; Subject: F(25,100) = 3.175, p<0.0001]. Statistical analysis utilized a two-way ANOVA with time point and group as factors and two-way RM ANOVA with time (seconds) and group as factors across 6- and 14-month-old mice. Tukey’s or Sidak’s post-hoc tests were performed where applicable. Error bars indicate SEM. p ≤ 0.05, **p ≤ 0.01, ***p ≤ 0.001, ****p ≤ 0.0001, ns = not significant. n= 8-11 per group after outlier removal.

### Negative engram stimulation increases glial cell number and changes morphology

We next tested the hypothesis that chronic negative engram activation will induce glial changes related to inflammation by performing cell counts and morphological characterization of both microglia (Iba1+) and astrocytes (GFAP+) in the hippocampus. For microglia (**Figure 5A-J**), 14-month-old mice in the hM3Dq group displayed an increase in Iba1+ cells, while 6-month-old mice did not (Two-way ANOVA; Interaction: F(1,8) = 7.615, p=0.0247; Age: F(1,8) = 3.259, p=0.1087; Group: F(1,8) = 13.42, p=0.0064)(Tukey’s [Group]: 6MO mCherry vs. hM3Dq, p=0.9906; 14MO mCherry vs. hM3Dq, p=0.0113. [Age] 6 vs. 14MO mCherry, p=0.9876; 6 vs. 14MO hM3Dq, p=0.0704)(**Figure 5C**). Additionally, these 14-month-old hM3Dq mice had microglia with increased ramification index, cell territory volume, minimum and maximum branch length, suggesting a distinct change in phenotype with negative memory activation (Two-way ANOVA; [Ramification index] Interaction: F(1,8) = 1.908, p=0.2046; Age: F(1,8) = 1.950, p=0.2001; Group: F(1,8) = 15.11, p=0.0046. Tukey’s [Group]: 6MO mCherry vs. hM3Dq, p=0.3513; 14MO mCherry vs. hM3Dq, p=0.03242 (**Figure 5D**). [Cell Territory Volume] Interaction: F(1,8) = 3.494, p=0.0965; Age: F(1,8) = 0.05044, p=0.8279; Group: F(1,8) = 16.45, p=0.0037. Tukey’s [Group]: 6MO mCherry vs. hM3Dq, p=0.4564; 14MO mCherry vs. hM3Dq, p=0.0129 (**Figure 5E**). [Min Branch Length] Interaction: F(1,8) = 8.287, p=0.0206; Age: F(1,8) = 9.619, p=0.0146; Group: F(1,8) = 7.470, p=0.0257. Tukey’s [Group]: 6MO mCherry vs. hM3Dq, p=0.9996; 14MO mCherry vs. hM3Dq, p=0.0174. [Age]: 6 vs. 14MO mCherry, p=0.9985; 6 vs. 14MO hM3Dq, p=0.0141 (**Figure 5I**). [Max Branch Length] Interaction: F(1,8) = 4.094, p=0.0777; Age: F(1,8) = 1.214, p=0.3026; Group: F(1,8) = 8.964, p=0.0172. Tukey’s [Group]: 6MO mCherry vs. hM3Dq, p=0.8994; 14MO mCherry vs. hM3Dq, p=0.0309 (**Figure 5J**). However, there were no significant differences in the average centroid distance, number of endpoints and average branch length for Iba1+ cells (Two-way ANOVA; [Avg Centroid Distance] Interaction: F(1,8) = 0.7707, p=0.4056; Age: F(1,8) = 1.155, p=0.3138; Group: F(1,8) = 5.459, p=0.0477. Tukey’s [Group]: 6MO mCherry vs. hM3Dq, p=0.7370; 14MO mCherry vs. hM3Dq, p=0.1838 (**Figure 5F**). [Number of Endpoints] Interaction: F(1,8) = 2.110, p=0.1884; Age: F(1,8) = 2.158, p=0.1801; Group: F(1,8) = 0.0237, p=0.8815 (**Figure 5G**). [Avg Branch Length] Interaction: F(1,8) = 0.7574, p=0.4095; Age: F(1,8) = 0.02109, p=0.8881; Group: F(1,8) = 0.1211, p=0.7369 (**Figure 5G**)).

**Figure 5:**
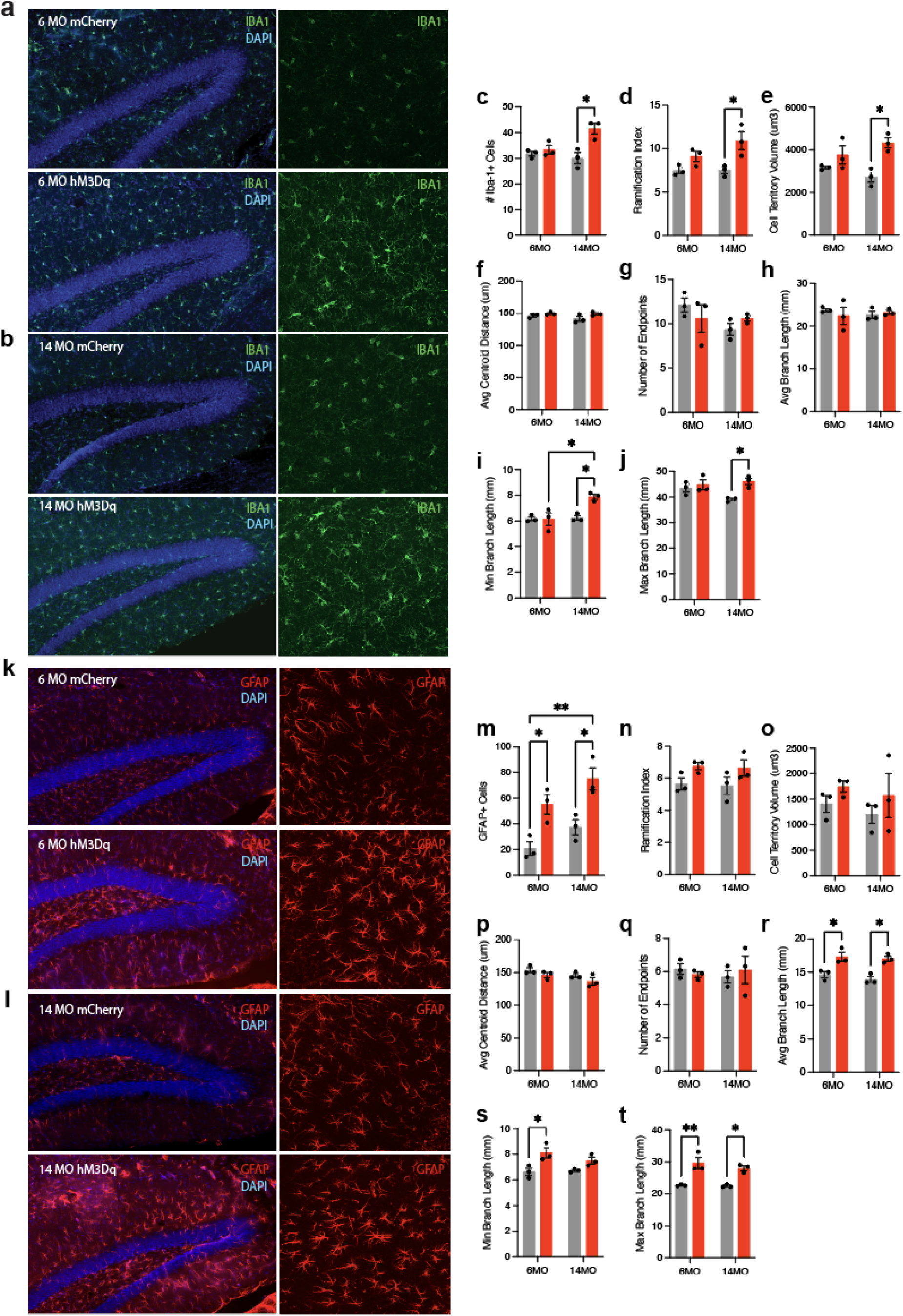
Chronic modulation of the negative engram induces changes in the number and morphology of microglia and astrocytes. (a-b) Representative images of Iba1+ microglia (green) in the hippocampus of (a) 6 month mCherry and hM3Dq mice and (b) 14 month mCherry and hM3Dq mice following 3 months of stimulation. (c) Number of Iba1+ cells. (d-j) Quantified morphological analysis of Iba1+ cells; (d) ramification index, (e) cell territory volume, (f) average centroid distance, (g) number of endpoints, (h) average branch length, (i) minimum branch length, and (j) maximum branch length. (k-l) Representative images of GFAP+ astrocytes (red) in the hippocampus of (k) 6 month mCherry and hM3Dq mice and (l) 14 month mCherry and hM3Dq mice following 3 months of stimulation. (m) Number of GFAP+ cells. (n-t) Quantified morphological analysis of GFAP+ cells; (n) ramification index, (o) cell territory volume, (p) average centroid distance, (q) number of endpoints, (r) average branch length, (s) minimum branch length, and (t) maximum branch length. Statistical analysis used a two-way ANOVA with time point and group as factors. Tukey’s post-hoc tests were performed. Error bars indicate SEM. p ≤ 0.05, **p ≤ 0.01, ***p ≤ 0.001, ****p ≤ 0.0001, ns = not significant. For cell counts, n=3 mice/group x 18 single tile ROIs each (GFAP, Iba1). For morphology, n=3 mice/group x 3 single tile ROIs each (GFAP, Iba1), each containing 20-30 glial cells.

For astrocytes (**Figure 5K-T**), there was a similar increase in the number of GFAP+ cells in the hippocampus, but in both the 6- and 14-month groups receiving hM3Dq activation (Two-way ANOVA; Interaction: F(1,8) = 0.05526, p=0.8201; Age: F(1,8) = 6.824, p=0.0310; Group: F(1,8) = 26.95, p=0.0008)(Tukey’s [Group]: 6MO mCherry vs. hM3Dq, p=0.0328; 14MO mCherry vs. hM3Dq, p=0.0208)(**Figure 5M**). Additionally, these astrocytes at both ages had increased average branch length and maximum branch length (Two-way ANOVA; [Avg Branch Length] Interaction: F(1,8) = 0.1537, p=0.7052; Age: F(1,8) = 0.9883, p=0.3493; Group: F(1,8) = 32.75, p=0.0004. Tukey’s [Group]: 6MO mCherry vs. hM3Dq, p=0.0228; 14MO mCherry vs. hM3Dq, p=0.0109 (**Figure 5R**). [Max Branch Length] Interaction: F(1,8) = 0.5438, p=0.4819; Age: F(1,8) = 0.8447, p=0.3849; Group: F(1,8) = 41.82, p=0.0002. Tukey’s [Group]: 6MO mCherry vs. hM3Dq, p=0.0041; 14MO mCherry vs. hM3Dq, p=0.0155 (**Figure 5T**)). Astrocytes in the 6-month hM3Dq group additionally had increased minimum branch length (Two-way ANOVA; Interaction: F(1,8) = 1.568, p=0.2459; Age: F(1,8) = 0.7927, p=0.3993; Group: F(1,8) = 16.26, p=0.0038)(Tukey’s [Group]: 6MO mCherry vs. hM3Dq, p=0.0238; 14MO mCherry vs. hM3Dq, p=0.2759)(**Figure 5S**). However, astrocytes did not show any significant differences for any group in their ramification index, cell territory volume, average centroid distance and number of endpoints (Two-way ANOVA; [Ramification Index] Interaction: F(1,8) = 0.0003, p=0.9856; Age: F(1,8) = 0.08582, p=0.7770; Group: F(1,8) = 6.677, p=0.0324. Tukey’s [Group]: 6MO mCherry vs. hM3Dq, p=0.3233; 14MO mCherry vs. hM3Dq, p=0.3339 (**Figure 5N**). [Cell Territory Volume] Interaction: F(1,8) = 0.0028, p=0.9584; Age: F(1,8) = 0.5819, p=0.4674; Group: F(1,8) = 1.996, p=0.1954 (**Figure 5O**). [Avg Centroid Distance] Interaction: F(1,8) = 0.04699, p=0.8338; Age: F(1,8) = 4.275, p=0.0725; Group: F(1,8) = 3.856, p=0.0852 (**Figure 5P**). [Number of Endpoints] Interaction: F(1,8) = 0.5463, p=0.4810; Age: F(1,8) = 0.0365, p=0.8532; Group: F(1,8) = 0.0071, p=0.9346)(**Figure 5Q**)). In summary, chronic activation of a negative engram induces an increase in glial cell number and changes their morphology in both young and older cohorts.

## Discussion

We chronically activated negative engrams in the vHPC using chemogenetics over the course of three months. This stimulation protocol induced detrimental behavioral effects as evidenced by increased anxiety-like behaviors, decreased working memory, and increased fear responses across extinction and generalization. Notably, our chronic negative engram activation significantly impacted microglial and astrocytic number and morphology within the hippocampus. Together, these findings present evidence for the role of negative memories in producing maladaptive cellular and behavioral responses within a rodent model and further our understanding of the broad impacts of chronic negative thinking-like behaviors that that may detrimentally affect human health.

### Molecular Features of Engram Cells

First, to characterize the molecular features of engram cells, we performed RNA-sequencing and utilized GSEA to compare gene expression profiles in cells that were labeled during a positive or negative experience (Rao-Ruiz et al., 2019; Fuentes-Ramos et al., 2021; Sardoo et al., 2022). Using *a priori* gene sets of interest, we found that negative engram cells upregulated genes that were associated with pathways involved in neurodegeneration, proinflammatory processes, and apoptosis. In contrast, positive engram cells upregulated genes involved in neuroprotective and anti-inflammatory processes. Notable genes that were significantly upregulated in the negative engram cells were secreted phosphoprotein 1 (*Spp1*) and transthyretin (*Ttr*). SPP1, also known as osteopontin, is an extracellular matrix protein widely expressed in immune cells, such as macrophages and microglia, mediating inflammatory responses. Blocking SPP1 protein expression in cells within the hippocampus has been shown to attenuate inflammation-induced depressive-like behaviors in mice (Li et al., 2023). Furthermore, the *Spp1* gene is consistently upregulated after chronic social stress (Stankiewicz et al., 2015), recall of contextual fear (Barnes, Kirtley & Thomas, 2012), and in rats with increased trait anxiety (Diaz-Moran et al., 2013). Mice receiving chronic negative memory stimulation in our study displayed similar increases in anxiety-like behaviors and impairments in fear responses. Further, TTR is a serum transporter for thyroid hormones and retinol binding proteins that has been shown to be detrimental when misfolded and aggregated in various forms of amyloidosis, while also having evidence of protective effects in Alzheimer’s Disease (AD) (Buxbaum et al., 2008; Sharma et al., 2019). Interestingly, TTR mRNA and protein were increased in both male and female rats that underwent acute or chronic stress, emphasizing the potential impact of glucocorticoid hormones on TTR protein levels (Martinho et al., 2012). Together, these findings suggest that genes upregulated in negative engram cells are linked to stress-induced changes in anxiety, fear and depressive-like behaviors.

Strikingly, negative engram cells also significantly upregulated amyloid precursor protein (*App*) and apolipoprotein E (*Apoe*) genes. APP plays a central role in AD pathology through generation of amyloid plaques (Lee et al., 2018; Lee et al., 2020). *Apoe* genotype accounts for the majority of AD risk and pathology (Johnson et al., 2015; Schuff et al., 2009), with the E4 isotype accounting for as much as 50% of AD in the United States (Raber et al., 2003). The upregulation of these genes involved in neurodegeneration and neuroinflammation is in line with human research showing a link between repetitive negative thinking and increased risk of cognitive decline and vulnerability to dementia (Schlosser et al., 2020). Furthermore, negative engram cells down-regulated neuroprotective and anti-inflammatory genes, notably brain derived neurotrophic factor (*Bdnf*). BDNF is a key molecule involved in memory in the healthy and pathological brain. Interventions such as exercise and antidepressant treatments enhance the expression of BDNF protein (Miranda et al., 2019). Future studies may further investigate the neuroprotective role of positive engrams to test, for instance, if activating a positive engram in a mouse model of AD confers protective effects on the brain and slows the progression of disease. Indeed, evidence of positive memory impact on mice with a depression-like phenotype has already shown promising results in this area of research (Ramirez et al., 2015). Together, these genetic data begin to uncover the molecular landscape that differentiates positive and negative memory-bearing cells, linking them to either protective or pathological brain states, respectively.

### Chronic stimulation of engrams alter behavior and cellular responses

Chronic modulation of engram cells in rodents has differential effects of behavior, depending on the brain region and valence of the tagged neurons. Reactivation of positive engram cells in the hippocampus rescued stress-induced depression-like phenotype and increased neurogenesis in mice (Ramirez et al., 2015). Further, chronic stimulation of dorsal or ventral hippocampal fear engrams produced a context-specific reduction or enhancement of fear responses, respectively (Chen et al., 2019). Interestingly, chronic optogenetic activation of dorsal hippocampal fear engrams results in context-specific reduction in fear expression that resembles extinction, even in mice with impaired extinction due to ethanol withdrawal (Cincotta et al., 2021). Activation of negative engram cells labeled during chronic social defeat stress induced social avoidance, a behavior common in human patients with MDD (Zhang et al., 2019). Together, these studies suggest that activity-dependent modulation of hippocampal cells over prolonged periods of time is sufficient to modify behavioral responses and brain states in mice.

Engram studies have predominantly focused on the neuronal phenotypes resulting from activity-dependent manipulations; however, glial cell responses to chronic memory activation and subsequent behavioral changes are unknown. Microglia are the brain’s resident immune cells, actively surveying the environment to maintain homeostasis and protect against insult and disease. Under normal conditions, microglial cells display small cell bodies and fine, highly ramified processes that allow them to detect and respond to threats (Nimmerjahn et al., 2005). Once a threat is detected (i.e. pathogens, debris, injury), microglia retract and exhibit large soma size and thickening, entering an ‘ameboid’ state. Evidence has accumulated that microglial morphology may be more complex than this dichotomy between ‘surveying’ and ‘ameboid’ microglia (Vidal-Itriago et al., 2022; Leyh et al., 2021; Reddaway et al., 2023). Our chronic negative engram activation may induce an ‘insult’ on the brain, as evidenced by an increase in Iba1 expression and hyper-ramified morphology, indicating a complex change in microglial phenotype. In support of these findings, chronic stress similarly induced an increase in Iba1 expression and hyper-ramified microglia in the prefrontal cortex (PFC) that was correlated with impaired spatial working memory (Hinwood et al., 2012). Further supporting these findings, a mouse model of PTSD displayed a similar increase in Iba1+ and hyper-ramified microglial cells in the PFC and CA1 layer of the hippocampus (Smith et al., 2019). Together, the changes in microglial cell number and morphology observed in the context of chronic negative engram stimulation are reminiscent of the alterations seen in chronic stress, PTSD and related depression-like behaviors.

Further, astrocytes express glial fibrillary acidic protein (GFAP), a critical intermediate filament responsible for maintaining their cytoskeletal structure (Yang and Wang, 2016). Chronic restraint stress has been associated with static GFAP+ astrocytic number and atrophy of astrocytic processes in the hippocampus, amygdala, and retrosplenial cortex (Kassem et al., 2013). Studies have primarily shown a decrease in GFAP+ astrocyte numbers and atrophy in rodent brains subjected to chronic and acute stressors (Tynan et al., 2013; Banasr and Duman, 2008; Gosselin et al., 2009; Miguel-Hidalgo et al., 2000; Bender, Calfa and Molina, 2016; Coldeluppi et al., 2021). However, recent studies have found a variety of changes in astrocytic morphology and number depending on the type of stressor and brain region. Conversely, others report an increase in GFAP+ cells in the hippocampus (Jang et al., 2008). Additionally, unpredictable chronic mild stress substantially increased GFAP immunoreactivity and promoted a more branched, reactive morphology in the hippocampus, which was correlated with increases in anxiety and anhedonic-like behaviors (Du Preez et al., 2021). This observed phenotype was coupled with similar changes in Iba1+ microglia, featuring increased branching complexity and ramification, longer processes and increased volume of occupation, as discussed previously (Du Preez et al., 2021; Hinwood et al., 2012). In the case of humans, the discourse on glial changes in chronic stress and depression-like behaviors has been similarly complex. GFAP concentrations in the cerebellum, prefrontal cortex, hippocampus and anterior cingulate cortex have been reported to decrease in patients with major depression compared to controls (Fatemi et al., 2003; Si et al., 2004; Webster et al., 2005; Cobb et al., 2015). However, a recent study found increased GFAP concentrations in the cerebral spinal fluid of patients with major depression compared to controls (Michel et al., 2021). Our findings that astrocytes increase in number and branch length in the hippocampus after chronic negative memory stimulation mirror the impacts of chronic stress on glial morphology. Further research is needed to understand these nuanced changes across brain regions, animal models, type of stressors and influence on behavior.

Our previous work demonstrated that global chronic activation of vHPC excitatory neurons or astrocytes for three months was sufficient to induce a variety of behavioral and cellular changes (Suthard et al., 2023). In our current study, we modulated hippocampal cells in an activity-dependent manner, gaining access to the specific impact of chronic negative memory activation on behavioral and cellular responses. Collectively, our data show a similar impact of CaMKII+ neuron and negative fear memory activation on fear extinction behaviors–namely, an increase in freezing responses. However, GFAP+ astrocyte activation induced a mild decrease in anxiety-related behaviors in the open field, while our negative memory activation significantly increased anxiety. Histologically, our previous work had mild impacts on glial number and morphology with non-memory-specific neuron or astrocyte chronic activation (Suthard et al., 2023). Our current study demonstrates a much more potent increase in both microglia and astrocyte number, as well as robust changes in morphological characteristics like ramification index, cell territory volume and branch lengths, which underscore the efficacy of chronic negative engram stimulation. A concern, however, is that activation of vHPC neurons could induce local tissue pathology as evidenced by inflammation and glial response. As this previous work activated all CaMKII+ neurons in vHPC and induced very modest changes in glial cells, we expect that in our current work, stimulating a smaller ensemble of cells, would induce less of such “off-target” inflammation. Together, these findings highlight the differences between modulating cells in a cell-type specific or activity-dependent manner (e.g. vHPC neurons or negative engram cells) and the resulting phenotypes that emerge.

Previous work from our lab and others has shown that there are differential effects of chronic activation of negative, positive and neutral ensembles within the hippocampus. As discussed earlier, chronic reactivation of a positive valence engram was sufficient to reverse the effects of stress on depression-like behaviors, whereas modulating a neutral ensemble did not (Ramirez et al, 2015). Here, we speculate that chronic activation of a positive engram in healthy, unstressed mice, may produce behavioral outcomes similar to those effects seen in chronic environmental enrichment, with improved cognitive flexibility, memory and even enhanced levels of anti-inflammatory proteins like BDNF (Bayne, 2018 *for review*). We further speculate that unlike neutral or negative memories, chronic stimulation of a positive memory may provide prophylactic effects against stress, degeneration, and aging, though this work remains an opportunity for future experiments. Other studies utilizing repeated negative memory activation (Chen et al, 2019; Zhang et al, 2019; Cincotta et al, 2021) showed content- and context-specific effects, such as bi-directional control over the strength of a fear memory in healthy and addiction-related states, with no changes observed in a neutral control group. Although our experiments here do not compare the effects of positive or neutral memory stimulation directly but rather compare a negative memory group to a fluorophore-only control, which is therefore a current limitation, we believe that chronic stimulation of each would produce unique sets of cellular and behavioral phenotypes that raise the intriguing notion of manipulating memories to mimic or ameliorate various pathological states.

### Relevance of chronic negative thinking-like behaviors to human health

Chronic memory activation may mimic symptoms of rumination, negative thinking and ongoing worry shared by individuals with GAD, MDD, and PTSD (Brewin et al., 2010; Newman et al., 2013; Watkins, 2008; McEvoy et al., 2013), but the mechanisms behind this connection are currently unknown. In the healthy brain, ‘inhibitory control’ of the prefrontal cortex on the hippocampus is thought to keep these symptoms at bay. Recent work has suggested a framework where stress may compromise GABAergic signaling in the hippocampus (Alaiyed et al., 2020; Banasr et al., 2017; Albrecht et al., 2021), allowing a state of hippocampal hyperactivity to persist. This elevated hippocampal activity and dysfunction of GABAergic interneuron is evident in human patients with major depression and PTSD, and the activity levels are correlated with behavioral rumination and flashback intensity (Osuch et al., 2001; Rayner, Jackson and Wilson, 2016). Human functional imaging work has supported this framework by showing that fronto-hippocampal inhibitory control via GABAergic signaling underlies this inability to suppress unwanted thoughts (Schmitz et al., 2017). Further, GABAergic signaling in the hippocampus has been shown to produce impaired extinction of conditioned fear (Muller et al., 2015) and increased anxiety (Crestani et al., 1999). This mirrors our findings of increased anxiety and decreased extinction ability in mice that experienced negative memory activation. Further exploration into how chronic engram activation impacts excitatory-inhibitory (E/I) imbalance may provide a link between memory modulation and induction of effects on the brain similar to those related to rumination, negative thinking, and worry.

In humans, persistent worrying and rumination behaviors that are common in neuropsychiatric disorders can affect the health of the entire body. In the context of our findings, the concept of allostatic load refers to the “wear and tear” the body experiences as a result of chronic exposure to stressful events. While the body attempts to produce a physiologically adaptive response through the process of allostasis, it is proposed that repeated cycles of allostasis in response to chronic stressors results in allostatic overload. In this state, the cumulative impact of chronic stress has overcome the body’s ability to cope, ultimately culminating in disease (McEwen and Stellar, 1993; McEwen, 2004). This applies to the nervous system, wherein rumination has been shown to increase the risk of alcohol and substance abuse, especially in young individuals, as a form of compensatory behavior to numb the persistent negative thoughts (Nolen-Hoeksema and Watkins, 2001). Rumination after a stressful event is also linked to insomnia, poor sleep quality and pre-sleep intrusive thoughts, as well as an increase in the risk of heart disease and cardiovascular issues via autonomic dysregulation (Guastella and Moulds, 2007; Busch, Possel, & Valentine, 2017). Together, we hope that the current study provides a novel framework in which memory itself may affect, and lead to, the maladaptive brain and body states described here.

### Chronic engram reactivation and its differing effects on freezing behavior

Previous work showed that experimental activation of recall-induced populations in the dentate gyrus leads to fear attenuation (Khalaf et al, 2018), while our experimental manipulation produces lasting negative impacts on the brain and behavior. Specifically, this study posits that this fear attenuation occurs in neuronal ensembles through updating or unlearning of the original memory trace via the engagement, rather than suppression, of an original traumatic experience. Notably, one major difference is that our experiment reactivated an ensemble that was tagged during memory encoding, while they activated a remote recall-induced ensemble that was tagged one month after encoding. Although there is high overlap between the encoding and recall ensembles when mice are exposed to the conditioning context, these ensembles are not identical and may result in different behavioral phenotypes when chronically reactivated. Further, Khalaf et al. relied on reactivation of the recall-induced ensemble during extinction to facilitate rapid fear attenuation. This differs experimentally from our current work, as their reactivation occurred during the extinction process in the previously conditioned context, while we reactivated chronically in the animal’s home cage over the course of a longer time period. It may be necessary that the memory is reactivated, and thus, more liable to re-contextualization, in the original context compared to an unrelated homecage environment where there are no related cues present. Importantly, this previous work tested the attenuation of fear shortly after an extinction process, while we did not extinguish the memory with aid of the memory reactivation. Finally, we tested remote recall (i.e. 3 months post-conditioning), as opposed to a shorter time interval (28 days). Despite these discrepancies, it is indeed possible that our chronic reactivation of a fear acquisition ensemble over time mildly attenuates or changes the original memory in some way. Future work could build on the findings of Khalaf et al. and/or show differences in the activation of a conditioning vs. recall-induced ensemble on fear attenuation during extinction.

### Limitations of the Study

#### Limitations of chronic chemogenetic modulation

In this study, we rely on the successful chemogenetic modulation of fearful neuronal ensembles to assess its impact on behavior over a long period of time. This strategy requires that our ligand, DCZ, maintains an effect on neuronal excitability across our three months of chronic reactivation. Previous work has shown that CNO administration in drinking water over one month consistently inhibited hM4Di+ neurons without altering baseline neuronal excitability as measured by firing rate and potassium currents (Xia et al, 2017). Although this is only for one month, it is administered via the same oral route as our DCZ protocol and suggests that at least for that amount of time we are likely producing similar effects. However, this must be experimentally tested as CNO and DCZ are unique ligands for chemogenetic modulation and must be extended out from one month to three months to assess the longevity of its effects in the brain. If DCZ is only having an effect for one vs. three months, we are still observing enduring changes that resulted from this shorter-term disturbance, which nonetheless informs our understanding of negative memory’s effects on the brain and behavior. On this same note, assessment of DREADD impact on E/I balance must be quantified utilizing both excitatory and inhibitory markers in future work to understand how chemogenetic modulation of engram cells may truly impact circuit-level E/I dysfunction.

Related to this, other work has shown evidence of DREADD toxicity at high titer levels of AAV2/7-CaMKII-hM4Di-mCherry in the HPC at five weeks (Goossens et al, 2021). We mitigated this risk by utilizing a viral strategy that targets a smaller number of engram cells using the AAV9-DIO-Flex-hM3Dq-mCherry virus at a lower titer, unlike this previous work that targeted all CaMKII+ cells in the HPC. Despite these efforts, we wanted to assess signs of decreasing neuronal health that may have resulted from chronic DREADD expression and utilized NeuN counts within multiple hippocampal subregions (Yousef et al, 2017). Here, we note that we did not observe significant decreases in NeuN across active Gq receptor or mCherry control groups in either of the age groups (Supplemental Figure 1). However, immunohistochemistry using an individual marker may not be sufficient to capture the entire health profile of an individual neuron and future work should consider other markers of cell death or inflammation to ensure prolonged health of DREADD+ cells.

#### Limitations of genetic tagging strategies

Further, this present study relies on use of the Tet-tag and TRAP2 genetic strategies to label engram cells for sequencing (Tet-tag) and chronic activation (TRAP2). While both systems rely on the *cFos* promoter to drive an effector of interest (fluorophore or DREADD), their efficacy and temporal resolution vary substantially depending on the genetic cell-type, brain region, temporal parameters of Dox or 4-OHT delivery, subject-by-subject metabolic variability and threshold to Dox induction given the promoter sequences inherent to each system.The TRAP2 line labels a subset of endogenously activeCA1 pyramidal cells (e.g. 5-18%) while the DOX system labels 20-40% of CA1 pyramidal cells (DeNardo et al, 2019; Monasterio et al, *BioRxiv* 2024). Further, the temporal windows for each system ranges from hours in TRAP2, compared to 24-48 hours in the Tet-tag system, resulting in potentially less-specific labeling in the latter (DeNardo et al, 2019; Denny et al, 2014; Liu & Ramirez et al, 2012)). Further, the efficacy of tagging a population of cells with either system will inherently constrain the number of cells that may overlap with cFos+ cells upon re-exposure to a given experience, given the difference in the percentage of initial cells labeled (Kim & Cho, 2020; Ortega-de San Luis et al, 2023; Shpokayte at al 2022). However, it is important to note that tagging vHPC cells with both strategies are nonetheless sufficient to drive behavioral responses (Shpokayte et al, 2022; Ortega-de San Luis et al, 2023).

Finally, and promisingly, as more studies continue to link the *in vivo* physiological dynamics of these cell populations tagged using each system (e.g. compare Pettit et al, 2022 with Tanaka et al, 2018) and correlating their activity to behavioral phenotypes, our field is in the prime position to uncover deeper principles governing hippocampus-mediated engrams in the brain. Together, we believe a more comprehensive understanding of these systems is fully warranted, especially in the service of further cataloging cellular similarities and differences within such tagged populations.

#### Limitations of gene set enrichment analysis within engram cells

Although the gene sets were pre-selected to assess changes canonically involved in “neurodegeneration” or “neuroprotection” such as *App, Apoe or Bdnf*, a major limitation of this technique is that we must avoid making strong claims about the actual function of these up- or down-regulated genes without performing proper causal studies. We hope these findings provide an unbiased inventory of the changes induced within engram cells related to memory valence that can inspire future work testing the knock-in or knock-down of these genes to assess their direct impact on the brain and behavior.

#### Limitations of immunohistochemistry analysis

Despite our study’s assessment of changes in astrocytic and microglial morphology and number after our manipulation, using immunohistochemistry does not allow us to capture all changes that may be occurring at the gene or protein level, painting an incomplete picture of the impact on glial cells. Future work utilizing transcriptomic or proteomic profiling of both engram and non-neuronal cells will help the field understand how these cell types are interacting with one another and provide a more complete mechanistic understanding of these glial-related changes.

## Acknowledgements

This material is based upon work supported by the Air Force Office of Scientific Research (AFOSR) under award number FA9550-21-1-0310, the Ludwig Family Foundation, an NIH Early Independence Award DP5 OD023106-01, NIH Transformative Award, Brain and Behavior Research Foundation Young Investigator Grant, McKnight Foundation Memory and Cognitive Disorders Award, Pew Scholars Program in the Biomedical Sciences, the Chan-Zuckerberg Initiative, and the Center for Systems Neuroscience and Neurophotonics Center at Boston University.

---

**Supplemental Figure 1:**
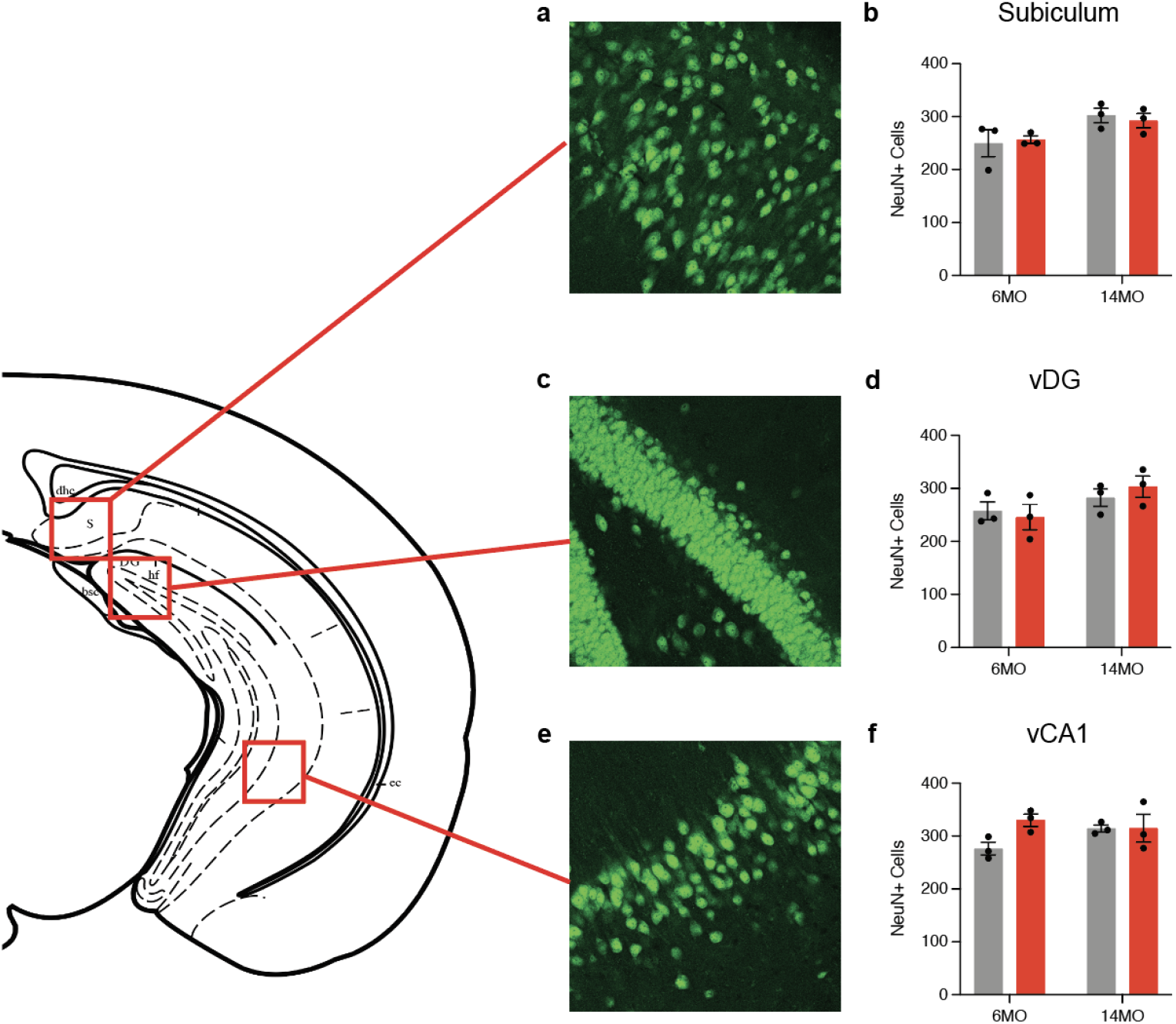
Chronic stimulation of the negative engram does not induce neuronal death. Representative images and quantification of NeuN+ cells in the (a-b) subiculum, (c-d) vDG, and (e-f) vCA1 of all groups (6 month mCherry and hM3Dq and 14 month mCherry and hM3Dq). Values are given as a mean + (SEM). Statistical analysis was performed with two-way ANOVA followed by Tukey’s post-hoc test. For cell counts, n=3 mice/group x 18 single tile ROIs each brain region (NeuN). No label = not significant, **p>0.001, ***p>0.0001, ****p>0.00001).

**Supplemental Figure 2:**
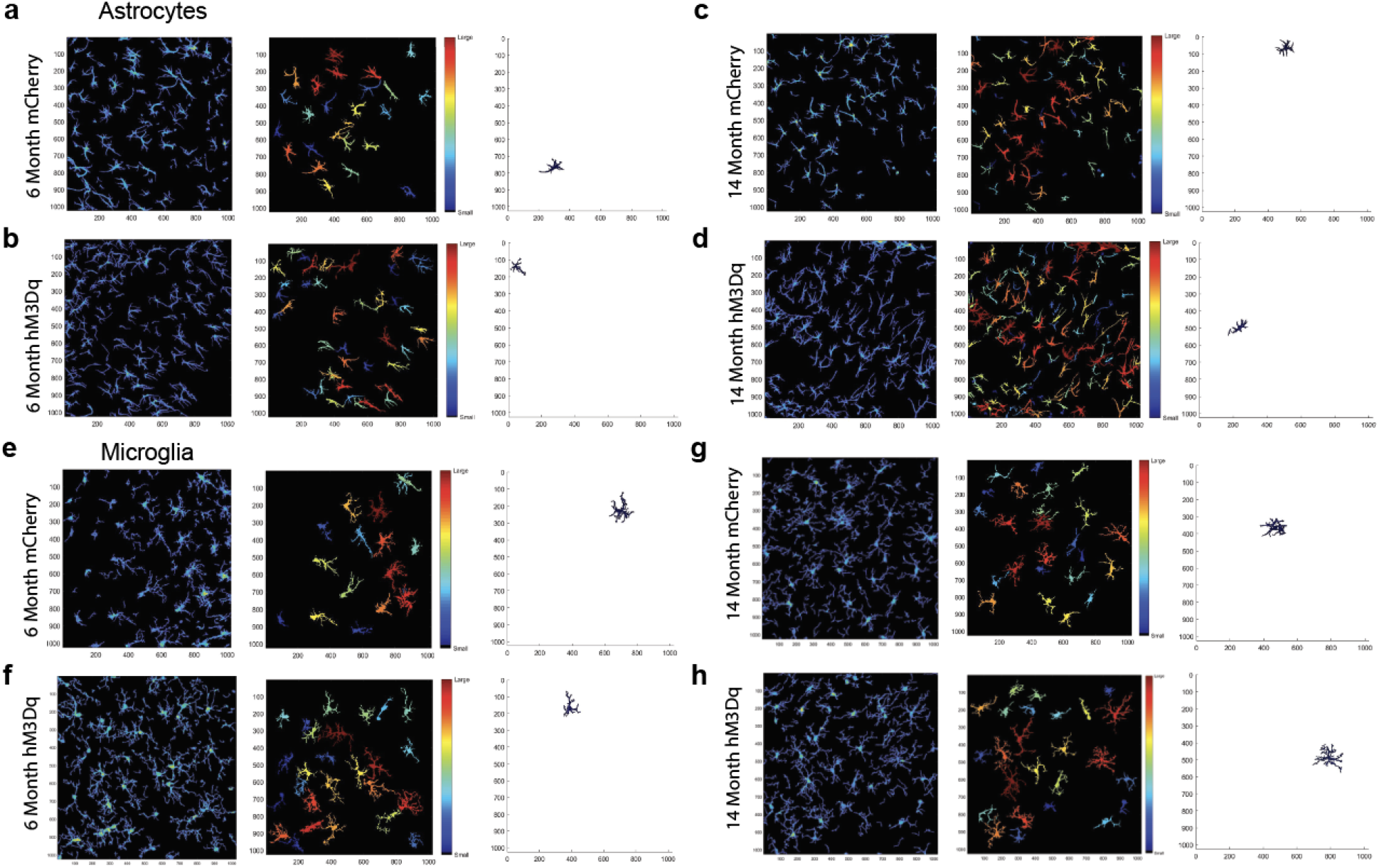
Analysis of astrocyte and microglial morphology using the MATLAB-based tool 3DMorph. a-d) Representative 3DMorph images of GFAP+ astrocytes in (a) 6 month mCherry, (b) 6 month hM3Dq, (c) 14 month mCherry, and (d) 14 month hM3Dq mice. (e-h) Representative 3DMorph images of Iba1+ microglia in (e) 6 month mCherry, (f) 6 month hM3Dq, (g) 14 month mCherry, and (h) 14 month hM3Dq mice.

## References

Alaiyed, S., McCann, M., Mahajan, G., Rajkowska, G., Stockmeier, C. A., Kellar, K. J., Wu, J. Y., & Conant, K. (2020). Venlafaxine Stimulates an MMP-9-Dependent Increase in Excitatory/Inhibitory Balance in a Stress Model of Depression. The Journal of neuroscience : the official journal of the Society for Neuroscience, 40(22), 4418–4431. 10.1523/JNEUROSCI.2387-19.2020

Albrecht, A., Redavide, E., Regev-Tsur, S., Stork, O., & Richter-Levin, G. (2021). Hippocampal GABAergic interneurons and their co-localized neuropeptides in stress vulnerability and resilience. Neuroscience and biobehavioral reviews, 122, 229–244. 10.1016/j.neubiorev.2020.11.002

Banasr, M., & Duman, R. S. (2008). Glial loss in the prefrontal cortex is sufficient to induce depressive-like behaviors. Biological psychiatry, 64(10), 863–870. 10.1016/j.biopsych.2008.06.008

Banasr, M., Lepack, A., Fee, C., Duric, V., Maldonado-Aviles, J., DiLeone, R., Sibille, E., Duman, R. S., & Sanacora, G. (2017). Characterization of GABAergic marker expression in the chronic unpredictable stress model of depression. Chronic stress (Thousand Oaks, Calif.), 1, 2470547017720459. 10.1177/2470547017720459

Barnes, P., Kirtley, A., & Thomas, K. L. (2012). Quantitatively and qualitatively different cellular processes are engaged in CA1 during the consolidation and reconsolidation of contextual fear memory. Hippocampus, 22(2), 149–171. 10.1002/hipo.20879

Bayne K. (2018). Environmental enrichment and mouse models: Current perspectives. Animal models and experimental medicine, 1(2), 82–90. 10.1002/ame2.12015

Bender, C. L., Calfa, G. D., & Molina, V. A. (2016). Astrocyte plasticity induced by emotional stress: A new partner in psychiatric physiopathology?. Progress in neuro-psychopharmacology & biological psychiatry, 65, 68–77. 10.1016/j.pnpbp.2015.08.005

Berg, S., Kutra, D., Kroeger, T., Straehle, C. N., Kausler, B. X., Haubold, C., Schiegg, M., Ales, J., Beier, T., Rudy, M., Eren, K., Cervantes, J. I., Xu, B., Beuttenmueller, F., Wolny, A., Zhang, C., Koethe, U., Hamprecht, F. A., & Kreshuk, A. (2019). ilastik: interactive machine learning for (bio)image analysis. Nature methods, 16(12), 1226–1232. 10.1038/s41592-019-0582-9

Brewin, C. R., Gregory, J. D., Lipton, M., & Burgess, N. (2010). Intrusive images in psychological disorders: characteristics, neural mechanisms, and treatment implications. Psychological review, 117(1), 210–232. 10.1037/a0018113

Busch, L. Y., Pössel, P., & Valentine, J. C. (2017). Meta-analyses of cardiovascular reactivity to rumination: A possible mechanism linking depression and hostility to cardiovascular disease. Psychological bulletin, 143(12), 1378–1394. 10.1037/bul0000119

Buxbaum, J. N., Ye, Z., Reixach, N., Friske, L., Levy, C., Das, P., Golde, T., Masliah, E., Roberts, A. R., & Bartfai, T. (2008). Transthyretin protects Alzheimer’s mice from the behavioral and biochemical effects of Abeta toxicity. Proceedings of the National Academy of Sciences of the United States of America, 105(7), 2681–2686. 10.1073/pnas.0712197105

Chen, B. K., Murawski, N. J., Cincotta, C., McKissick, O., Finkelstein, A., Hamidi, A. B., Merfeld, E., Doucette, E., Grella, S. L., Shpokayte, M., Zaki, Y., Fortin, A., & Ramirez, S. (2019). Artificially Enhancing and Suppressing Hippocampus-Mediated Memories. Current biology : CB, 29(11), 1885–1894.e4. 10.1016/j.cub.2019.04.065

Cobb, J. A., O’Neill, K., Milner, J., Mahajan, G. J., Lawrence, T. J., May, W. L., Miguel-Hidalgo, J., Rajkowska, G., & Stockmeier, C. A. (2016). Density of GFAP-immunoreactive astrocytes is decreased in left hippocampi in major depressive disorder. Neuroscience, 316, 209–220. 10.1016/j.neuroscience.2015.12.044

Codeluppi, S. A., Chatterjee, D., Prevot, T. D., Bansal, Y., Misquitta, K. A., Sibille, E., & Banasr, M. (2021). Chronic Stress Alters Astrocyte Morphology in Mouse Prefrontal Cortex. The international journal of neuropsychopharmacology, 24(10), 842–853. 10.1093/ijnp/pyab052

Cincotta, C., Murawski, N. J., Grella, S. L., McKissick, O., Doucette, E., & Ramirez, S. (2021). Chronic activation of fear engrams induces extinction-like behavior in ethanol-exposed mice. Hippocampus, 31(1), 3–10. 10.1002/hipo.23263

Crestani, F., Lorez, M., Baer, K., Essrich, C., Benke, D., Laurent, J. P., Belzung, C., Fritschy, J. M., Lüscher, B., & Mohler, H. (1999). Decreased GABAA-receptor clustering results in enhanced anxiety and a bias for threat cues. Nature neuroscience, 2(9), 833–839. 10.1038/12207

DeNardo, L. A., Liu, C. D., Allen, W. E., Adams, E. L., Friedmann, D., Fu, L., Guenthner, C. J., Tessier-Lavigne, M., & Luo, L. (2019). Temporal evolution of cortical ensembles promoting remote memory retrieval. Nature neuroscience, 22(3), 460–469. 10.1038/s41593-018-0318-7

Denny, C. A., Kheirbek, M. A., Alba, E. L., Tanaka, K. F., Brachman, R. A., Laughman, K. B., Tomm, N. K., Turi, G. F., Losonczy, A., & Hen, R. (2014). Hippocampal memory traces are differentially modulated by experience, time, and adult neurogenesis. Neuron, 83(1), 189–201. 10.1016/j.neuron.2014.05.018

Dobin, A., Davis, C. A., Schlesinger, F., Drenkow, J., Zaleski, C., Jha, S., Batut, P., Chaisson, M., & Gingeras, T. R. (2013). STAR: ultrafast universal RNA-seq aligner. Bioinformatics (Oxford, England), 29(1), 15–21. 10.1093/bioinformatics/bts635

Díaz-Morán, S., Palència, M., Mont-Cardona, C., Cañete, T., Blázquez, G., Martínez-Membrives, E., López-Aumatell, R., Sabariego, M., Donaire, R., Morón, I., Torres, C., Martínez-Conejero, J. A., Tobeña, A., Esteban, F. J., & Fernández-Teruel, A. (2013). Gene expression in the hippocampus as a function of differential trait anxiety levels in genetically heterogeneous NIH-HS rats. Behavioral brain research, 257, 129–139. 10.1016/j.bbr.2013.09.041

Du Preez, A., Onorato, D., Eiben, I., Musaelyan, K., Egeland, M., Zunszain, P. A., Fernandes, C., Thuret, S., & Pariante, C. M. (2021). Chronic stress followed by social isolation promotes depressive-like behavior, alters microglial and astrocyte biology and reduces hippocampal neurogenesis in male mice. Brain, behavior, and immunity, 91, 24–47. 10.1016/j.bbi.2020.07.015

Fatemi, S. H., Laurence, J. A., Araghi-Niknam, M., Stary, J. M., Schulz, S. C., Lee, S., & Gottesman, I. I. (2004). Glial fibrillary acidic protein is reduced in the cerebellum of subjects with major depression, but not schizophrenia. Schizophrenia research, 69(2-3), 317–323. 10.1016/j.schres.2003.08.014

Ferrari, L.L., Ogbeide-Latario, O.E., Gompf, H.S., Anaclet, C., 2022. Validation of DREADD agonists and administration route in a murine model of sleep enhancement. J Neurosci Methods 380. 10.1016/J.JNEUMETH.2022.109679

Fuentes-Ramos, M., Alaiz-Noya, M., & Barco, A. (2021). Transcriptome and epigenome analysis of engram cells: Next-generation sequencing technologies in memory research. Neuroscience and biobehavioral reviews, 127, 865–875. 10.1016/j.neubiorev.2021.06.010

Goossens, M. G., Larsen, L. E., Vergaelen, M., Wadman, W., Van den Haute, C., Brackx, W., Proesmans, S., Desloovere, J., Christiaen, E., Craey, E., Vanhove, C., Vonck, K., Boon, P., & Raedt, R. (2021). Level of hM4D(Gi) DREADD Expression Determines Inhibitory and Neurotoxic Effects in the Hippocampus. eNeuro, 8(6), ENEURO.0105-21.2021. 10.1523/ENEURO.0105-21.2021

Gosselin, R. D., Gibney, S., O’Malley, D., Dinan, T. G., & Cryan, J. F. (2009). Region specific decrease in glial fibrillary acidic protein immunoreactivity in the brain of a rat model of depression. Neuroscience, 159(2), 915–925. 10.1016/j.neuroscience.2008.10.018

Guastella, A. J., & Moulds, M. L. (2007). The impact of rumination on sleep quality following a stressful life event. Personality and Individual Differences, 42(6), 1151–1162. 10.1016/j.paid.2006.04.028

Hinwood, M., Tynan, R. J., Charnley, J. L., Beynon, S. B., Day, T. A., & Walker, F. R. (2013). Chronic stress induced remodeling of the prefrontal cortex: structural re-organization of microglia and the inhibitory effect of minocycline. Cerebral cortex (New York, N.Y. : 1991), 23(8), 1784–1797. 10.1093/cercor/bhs151

Jang, S., Suh, S. H., Yoo, H. S., Lee, Y. M., & Oh, S. (2008). Changes in iNOS, GFAP and NR1 expression in various brain regions and elevation of sphingosine-1-phosphate in serum after immobilized stress. Neurochemical research, 33(5), 842–851. 10.1007/s11064-007-9523-6

Johnson, L. A., Zuloaga, D. G., Bidiman, E., Marzulla, T., Weber, S., Wahbeh, H., & Raber, J. (2015). ApoE2 Exaggerates PTSD-Related Behavioral, Cognitive, and Neuroendocrine Alterations. Neuropsychopharmacology : official publication of the American College of Neuropsychopharmacology, 40(10), 2443–2453. 10.1038/npp.2015.95

Kassem, M. S., Lagopoulos, J., Stait-Gardner, T., Price, W. S., Chohan, T. W., Arnold, J. C., Hatton, S. N., & Bennett, M. R. (2013). Stress-induced grey matter loss determined by MRI is primarily due to loss of dendrites and their synapses. Molecular neurobiology, 47(2), 645–661. 10.1007/s12035-012-8365-7

Khalaf, O., Resch, S., Dixsaut, L., Gorden, V., Glauser, L., & Gräff, J. (2018). Reactivation of recall-induced neurons contributes to remote fear memory attenuation. Science (New York, N.Y.), 360(6394), 1239–1242. 10.1126/science.aas9875

Kim, W. B., & Cho, J. H. (2020). Encoding of contextual fear memory in hippocampal-amygdala circuit. Nature communications, 11(1), 1382. 10.1038/s41467-020-15121-2

Lee, M. H., Siddoway, B., Kaeser, G. E., Segota, I., Rivera, R., Romanow, W. J., Liu, C. S., Park, C., Kennedy, G., Long, T., & Chun, J. (2018). Somatic APP gene recombination in Alzheimer’s disease and normal neurons. Nature, 563(7733), 639–645. 10.1038/s41586-018-0718-6

Lee, S. H., Kang, J., Ho, A., Watanabe, H., Bolshakov, V. Y., & Shen, J. (2020). APP Family Regulates Neuronal Excitability and Synaptic Plasticity but Not Neuronal Survival. Neuron, 108(4), 676–690.e8. 10.1016/j.neuron.2020.08.011

Leyh, J., Paeschke, S., Mages, B., Michalski, D., Nowicki, M., Bechmann, I., & Winter, K. (2021). Classification of Microglial Morphological Phenotypes Using Machine Learning. Frontiers in cellular neuroscience, 15, 701673. 10.3389/fncel.2021.701673

Li, T., Yuan, L., Zhao, Y., Jiang, Z., Gai, C., Xin, D., Ke, H., Guo, X., Chen, W., Liu, D., Wang, Z., & Ho, C. S. H. (2023). Blocking osteopontin expression attenuates neuroinflammation and mitigates LPS-induced depressive-like behavior in mice. Journal of affective disorders, 330, 83–93. 10.1016/j.jad.2023.02.105

Liao, Y., Smyth, G. K., & Shi, W. (2014). featureCounts: an efficient general purpose program for assigning sequence reads to genomic features. Bioinformatics (Oxford, England), 30(7), 923–930. 10.1093/bioinformatics/btt656

Liu, X., Ramirez, S., Pang, P. T., Puryear, C. B., Govindarajan, A., Deisseroth, K., & Tonegawa, S. (2012). Optogenetic stimulation of a hippocampal engram activates fear memory recall. Nature, 484(7394), 381–385. 10.1038/nature11028

Love, M. I., Huber, W., & Anders, S. (2014). Moderated estimation of fold change and dispersion for RNA-seq data with DESeq2. Genome biology, 15(12), 550. 10.1186/s13059-014-0550-8

Marchant, N. L., Lovland, L. R., Jones, R., Pichet Binette, A., Gonneaud, J., Arenaza-Urquijo, E. M., Chételat, G., Villeneuve, S., & PREVENT-AD Research Group (2020). Repetitive negative thinking is associated with amyloid, tau, and cognitive decline. Alzheimer’s & dementia : the journal of the Alzheimer’s Association, 16(7), 1054–1064. 10.1002/alz.12116

Maren, S., & Holmes, A. (2016). Stress and Fear Extinction. Neuropsychopharmacology : official publication of the American College of Neuropsychopharmacology, 41(1), 58–79. 10.1038/npp.2015.180

Martinho, A., Gonçalves, I., Costa, M., & Santos, C. R. (2012). Stress and glucocorticoids increase transthyretin expression in rat choroid plexus via mineralocorticoid and glucocorticoid receptors. Journal of molecular neuroscience : MN, 48(1), 1–13. 10.1007/s12031-012-9715-7

McEvoy, P. M., Watson, H., Watkins, E. R., & Nathan, P. (2013). The relationship between worry, rumination, and comorbidity: evidence for repetitive negative thinking as a transdiagnostic construct. Journal of affective disorders, 151(1), 313–320. 10.1016/j.jad.2013.06.014

McEwen, B. S., & Stellar, E. (1993). Stress and the individual. Mechanisms leading to disease. Archives of internal medicine, 153(18), 2093–2101.

McEwen B. S. (2004). Protection and damage from acute and chronic stress: allostasis and allostatic overload and relevance to the pathophysiology of psychiatric disorders. Annals of the New York Academy of Sciences, 1032, 1–7. 10.1196/annals.1314.001

McLaughlin, K. A., Conron, K. J., Koenen, K. C., & Gilman, S. E. (2010). Childhood adversity, adult stressful life events, and risk of past-year psychiatric disorder: a test of the stress sensitization hypothesis in a population-based sample of adults. Psychological medicine, 40(10), 1647–1658. 10.1017/S0033291709992121

Michel, M., Fiebich, B. L., Kuzior, H., Meixensberger, S., Berger, B., Maier, S., Nickel, K., Runge, K., Denzel, D., Pankratz, B., Schiele, M. A., Domschke, K., van Elst, L. T., & Endres, D. (2021). Increased GFAP concentrations in the cerebrospinal fluid of patients with unipolar depression. Translational psychiatry, 11(1), 308. 10.1038/s41398-021-01423-6

Miguel-Hidalgo, J. J., Baucom, C., Dilley, G., Overholser, J. C., Meltzer, H. Y., Stockmeier, C. A., & Rajkowska, G. (2000). Glial fibrillary acidic protein immunoreactivity in the prefrontal cortex distinguishes younger from older adults in major depressive disorder. Biological psychiatry, 48(8), 861–873. 10.1016/s0006-3223(00)00999-9

Miranda, M., Morici, J. F., Zanoni, M. B., & Bekinschtein, P. (2019). Brain-Derived Neurotrophic Factor: A Key Molecule for Memory in the Healthy and the Pathological Brain. Frontiers in cellular neuroscience, 13, 363. 10.3389/fncel.2019.00363

Monasterio, A., Lienkaemper, C., Coello, S., Ocker, G.K., Ramirez, S., Scott, B.B. (2024) CA1 engram cell dynamics before and after learning. bioRxiv, 10.1101/2024.04.16.589790

Mootha, V. K., Lindgren, C. M., Eriksson, K. F., Subramanian, A., Sihag, S., Lehar, J., Puigserver, P., Carlsson, E., Ridderstråle, M., Laurila, E., Houstis, N., Daly, M. J., Patterson, N., Mesirov, J. P., Golub, T. R., Tamayo, P., Spiegelman, B., Lander, E. S., Hirschhorn, J. N., Altshuler, D., … Groop, L. C. (2003). PGC-1alpha-responsive genes involved in oxidative phosphorylation are coordinately downregulated in human diabetes. Nature genetics, 34(3), 267–273. 10.1038/ng1180

Müller, I., Çalışkan, G., & Stork, O. (2015). The GAD65 knock out mouse - a model for GABAergic processes in fear- and stress-induced psychopathology. Genes, brain, and behavior, 14(1), 37–45. 10.1111/gbb.12188

Nagai, Y., Miyakawa, N., Takuwa, H., Hori, Y., Oyama, K., Ji, B., Takahashi, M., Huang, X. P., Slocum, S. T., DiBerto, J. F., Xiong, Y., Urushihata, T., Hirabayashi, T., Fujimoto, A., Mimura, K., English, J. G., Liu, J., Inoue, K. I., Kumata, K., Seki, C., … Minamimoto, T. (2020). Deschloroclozapine, a potent and selective chemogenetic actuator enables rapid neuronal and behavioral modulations in mice and monkeys. Nature neuroscience, 23(9), 1157–1167. 10.1038/s41593-020-0661-3

Newman, M. G., Llera, S. J., Erickson, T. M., Przeworski, A., & Castonguay, L. G. (2013). Worry and generalized anxiety disorder: a review and theoretical synthesis of evidence on nature, etiology, mechanisms, and treatment. Annual review of clinical psychology, 9, 275–297. 10.1146/annurev-clinpsy-050212-185544

Nolen-Hoeksema, S., & Watkins, E. R. (2011). A Heuristic for Developing Transdiagnostic Models of Psychopathology: Explaining Multifinality and Divergent Trajectories. Perspectives on Psychological Science, 6(6), 589–609. 10.1177/1745691611419672

Nimmerjahn, A., Kirchhoff, F., & Helmchen, F. (2005). Resting microglial cells are highly dynamic surveillants of brain parenchyma in vivo. Science (New York, N.Y.), 308(5726), 1314–1318. 10.1126/science.1110647

Ortega-de San Luis, C., Pezzoli, M., Urrieta, E., & Ryan, T. J. (2023). Engram cell connectivity as a mechanism for information encoding and memory function. Current biology : CB, 33(24), 5368–5380.e5. 10.1016/j.cub.2023.10.074

Osuch, E. A., Benson, B., Geraci, M., Podell, D., Herscovitch, P., McCann, U. D., & Post, R. M. (2001). Regional cerebral blood flow correlated with flashback intensity in patients with posttraumatic stress disorder. Biological psychiatry, 50(4), 246–253. 10.1016/s0006-3223(01)01107-6

Pettit, N. L., Yap, E. L., Greenberg, M. E., & Harvey, C. D. (2022). Fos ensembles encode and shape stable spatial maps in the hippocampus. Nature, 609(7926), 327–334. 10.1038/s41586-022-05113-1

Raber, J., Huang, Y., & Ashford, J. W. (2004). ApoE genotype accounts for the vast majority of AD risk and AD pathology. Neurobiology of aging, 25(5), 641–650. 10.1016/j.neurobiolaging.2003.12.023

Ramirez, S., Liu, X., MacDonald, C. J., Moffa, A., Zhou, J., Redondo, R. L., & Tonegawa, S. (2015). Activating positive memory engrams suppresses depression-like behaviour. Nature, 522(7556), 335–339. 10.1038/nature14514

Ramirez, S., Tonegawa, S., & Liu, X. (2014). Identification and optogenetic manipulation of memory engrams in the hippocampus. Frontiers in behavioral neuroscience, 7, 226. 10.3389/fnbeh.2013.00226

Rao-Ruiz, P., Couey, J.J., Marcelo, I.M. et al. Engram-specific transcriptome profiling of contextual memory consolidation. Nat Commun 10, 2232 (2019). 10.1038/s41467-019-09960-x

Rayner, G., Jackson, G., & Wilson, S. (2016). Cognition-related brain networks underpin the symptoms of unipolar depression: Evidence from a systematic review. Neuroscience and biobehavioral reviews, 61, 53–65. 10.1016/j.neubiorev.2015.09.022

Reddaway, J., Richardson, P. E., Bevan, R. J., Stoneman, J., & Palombo, M. (2023). Microglial morphometric analysis: so many options, so little consistency. Frontiers in neuroinformatics, 17, 1211188. 10.3389/fninf.2023.1211188

Sardoo, A. M., Zhang, S., Ferraro, T. N., Keck, T. M., & Chen, Y. (2022). Decoding brain memory formation by single-cell RNA sequencing. Briefings in bioinformatics, 23(6), bbac412. 10.1093/bib/bbac412

Schlosser, M., Demnitz-King, H., Whitfield, T., Wirth, M., & Marchant, N. L. (2020). Repetitive negative thinking is associated with subjective cognitive decline in older adults: a cross-sectional study. BMC psychiatry, 20(1), 500. 10.1186/s12888-020-02884-7

Schuff, N., Woerner, N., Boreta, L., Kornfield, T., Shaw, L. M., Trojanowski, J. Q., Thompson, P. M., Jack, C. R., Jr, Weiner, M. W., & Alzheimer’s Disease Neuroimaging Initiative (2009). MRI of hippocampal volume loss in early Alzheimer’s disease in relation to ApoE genotype and biomarkers. Brain : a journal of neurology, 132(Pt 4), 1067–1077. 10.1093/brain/awp007

Shannon, P., Markiel, A., Ozier, O., Baliga, N. S., Wang, J. T., Ramage, D., Amin, N., Schwikowski, B., & Ideker, T. (2003). Cytoscape: a software environment for integrated models of biomolecular interaction networks. Genome research, 13(11), 2498–2504. 10.1101/gr.1239303

Sharma, M., Khan, S., Rahman, S., & Singh, L. R. (2019). The Extracellular Protein, Transthyretin Is an Oxidative Stress Biomarker. Frontiers in physiology, 10, 5. 10.3389/fphys.2019.00005

Shpokayte, M., McKissick, O., Guan, X., Yuan, B., Rahsepar, B., Fernandez, F. R., Ruesch, E., Grella, S. L., White, J. A., Liu, X. S., & Ramirez, S. (2022). Hippocampal cells segregate positive and negative engrams. Communications biology, 5(1), 1009. 10.1038/s42003-022-03906-8

Si, X., Miguel-Hidalgo, J. J., O’Dwyer, G., Stockmeier, C. A., & Rajkowska, G. (2004). Age-dependent reductions in the level of glial fibrillary acidic protein in the prefrontal cortex in major depression. Neuropsychopharmacology : official publication of the American College of Neuropsychopharmacology, 29(11), 2088–2096. 10.1038/sj.npp.1300525

Smith, K. L., Kassem, M. S., Clarke, D. J., Kuligowski, M. P., Bedoya-Pérez, M. A., Todd, S. M., Lagopoulos, J., Bennett, M. R., & Arnold, J. C. (2019). Microglial cell hyper-ramification and neuronal dendritic spine loss in the hippocampus and medial prefrontal cortex in a mouse model of PTSD. Brain, behavior, and immunity, 80, 889–899. 10.1016/j.bbi.2019.05.042

Stankiewicz, A. M., Goscik, J., Majewska, A., Swiergiel, A. H., & Juszczak, G. R. (2015). The Effect of Acute and Chronic Social Stress on the Hippocampal Transcriptome in Mice. PloS one, 10(11), e0142195. 10.1371/journal.pone.0142195

Subramanian, A., Tamayo, P., Mootha, V. K., Mukherjee, S., Ebert, B. L., Gillette, M. A., Paulovich, A., Pomeroy, S. L., Golub, T. R., Lander, E. S., & Mesirov, J. P. (2005). Gene set enrichment analysis: a knowledge-based approach for interpreting genome-wide expression profiles. Proceedings of the National Academy of Sciences of the United States of America, 102(43), 15545–15550. 10.1073/pnas.0506580102

Suthard, R. L., Jellinger, A. L., Surets, M., Shpokayte, M., Pyo, A. Y., Buzharsky, M. D., Senne, R. A., Dorst, K., Leblanc, H., & Ramirez, S. (2023). Chronic Gq activation of ventral hippocampal neurons and astrocytes differentially affects memory and behavior. Neurobiology of aging, 125, 9–31. 10.1016/j.neurobiolaging.2023.01.007

Tanaka, K. Z., He, H., Tomar, A., Niisato, K., Huang, A. J. Y., & McHugh, T. J. (2018). The hippocampal engram maps experience but not place. Science (New York, N.Y.), 361(6400), 392–397. 10.1126/science.aat5397

Tran, I., & Gellner, A. K. (2023). Long-term effects of chronic stress models in adult mice. Journal of neural transmission (Vienna, Austria : 1996), 130(9), 1133–1151. 10.1007/s00702-023-02598-6

Tynan, R. J., Beynon, S. B., Hinwood, M., Johnson, S. J., Nilsson, M., Woods, J. J., & Walker, F. R. (2013). Chronic stress-induced disruption of the astrocyte network is driven by structural atrophy and not loss of astrocytes. Acta neuropathologica, 126(1), 75–91. 10.1007/s00401-013-1102-0

Vidal-Itriago, A., Radford, R. A. W., Aramideh, J. A., Maurel, C., Scherer, N. M., Don, E. K., Lee, A., Chung, R. S., Graeber, M. B., & Morsch, M. (2022). Microglia morphophysiological diversity and its implications for the CNS. Frontiers in immunology, 13, 997786. 10.3389/fimmu.2022.997786

Wang, W., Qin, X., Wang, R., Xu, J., Wu, H., Khalid, A., Jiang, H., Liu, D., & Pan, F. (2020). EZH2 is involved in vulnerability to neuroinflammation and depression-like behaviors induced by chronic stress in different aged mice. Journal of affective disorders, 272, 452–464. 10.1016/j.jad.2020.03.154

Warde-Farley, D., Donaldson, S. L., Comes, O., Zuberi, K., Badrawi, R., Chao, P., Franz, M., Grouios, C., Kazi, F., Lopes, C. T., Maitland, A., Mostafavi, S., Montojo, J., Shao, Q., Wright, G., Bader, G. D., & Morris, Q. (2010). The GeneMANIA prediction server: biological network integration for gene prioritization and predicting gene function. Nucleic acids research, 38(Web Server issue), W214–W220. 10.1093/nar/gkq537

Watkins E. R. (2008). Constructive and unconstructive repetitive thought. Psychological bulletin, 134(2), 163–206. 10.1037/0033-2909.134.2.163

Webster, M. J., O’Grady, J., Kleinman, J. E., & Weickert, C. S. (2005). Glial fibrillary acidic protein mRNA levels in the cingulate cortex of individuals with depression, bipolar disorder and schizophrenia. Neuroscience, 133(2), 453–461. 10.1016/j.neuroscience.2005.02.037

Xia, F., Richards, B. A., Tran, M. M., Josselyn, S. A., Takehara-Nishiuchi, K., & Frankland, P. W. (2017). Parvalbumin-positive interneurons mediate neocortical-hippocampal interactions that are necessary for memory consolidation. eLife, 6, e27868. 10.7554/eLife.27868

Yang, Z., & Wang, K. K. (2015). Glial fibrillary acidic protein: from intermediate filament assembly and gliosis to neurobiomarker. Trends in neurosciences, 38(6), 364–374. 10.1016/j.tins.2015.04.003

York, E. M., LeDue, J. M., Bernier, L. P., & MacVicar, B. A. (2018). 3DMorph Automatic Analysis of Microglial Morphology in Three Dimensions from Ex Vivo and In Vivo Imaging. eNeuro, 5(6), ENEURO.0266-18.2018. 10.1523/ENEURO.0266-18.2018

Yousef, A., Robinson, J. L., Irwin, D. J., Byrne, M. D., Kwong, L. K., Lee, E. B., Xu, Y., Xie, S. X., Rennert, L., Suh, E., Van Deerlin, V. M., Grossman, M., Lee, V. M., & Trojanowski, J. Q. (2017). Neuron loss and degeneration in the progression of TDP-43 in frontotemporal lobar degeneration. Acta neuropathologica communications, 5(1), 68. 10.1186/s40478-017-0471-3

Zhan, J., Komal, R., Keenan, W.T., Hattar, S., Fernandez, D.C., 2019. Non-invasive Strategies for Chronic Manipulation of DREADD-controlled Neuronal Activity. J Vis Exp 2019. 10.3791/59439

Zhang, T. R., Larosa, A., Di Raddo, M. E., Wong, V., Wong, A. S., & Wong, T. P. (2019). Negative Memory Engrams in the Hippocampus Enhance the Susceptibility to Chronic Social Defeat Stress. The Journal of neuroscience : the official journal of the Society for Neuroscience, 39(38), 7576–7590. 10.1523/JNEUROSCI.1958-18.2019

